# Binding and activation of serotonergic G-protein coupled receptors by the multimodal antidepressant vortioxetine

**DOI:** 10.1101/2021.01.05.425370

**Authors:** Lucy Kate Ladefoged, Rebekka Koch, Philip C. Biggin, Birgit Schiøtt

**Affiliations:** Department of Chemistry, Aarhus University, Langelandsgade 140, 8000 Aarhus C, Denmark; Department of Biochemistry, South Parks Road, Oxford OX1 3QU, United Kingdom; Interdisciplinary Nanoscience Center (iNANO), Aarhus University, Gustav Wieds Vej 14, 8000 Aarhus C, Denmark

## Abstract

G-protein coupled receptors are important pharmacological targets. Despite substantial progress, important questions still remain concerning the details of activation: how can a ligand act as an agonist in one receptor, but as an antagonist in a homologous receptor, and how can agonists activate a receptor despite lacking polar functional groups able to interact with helix 5? Studying vortioxetine, an important multimodal antidepressant drug, may elucidate both questions. Herein, we present a thorough *in silico* analysis of vortioxetine binding to 5-HT_1A_, 5-HT_1B_, and 5-HT_7_ receptors and compare to available experimental data. We are able to rationalize the differential mode of action of vortioxetine at different receptors, but also, in the case of the 5-HT_1A_ receptor, we observe the initial steps of activation suggesting that interaction with helix 5 does not necessarily require a hydrogen bond as previously suggested. The results extend our current understanding of agonist and antagonist action at GPCRs.

## Introduction

G-protein coupled receptors (GPCRs) are major pharmacological targets: GPCRs constitute 12 % of the human genome and approximately 33 % of approved pharmacotherapies target GPCRs (Santos et al., 2017). Many aspects of the ligand binding process and activation mechanism have been elucidated (Figure 1, (Latorraca, Venkatakrishnan, & Dror, 2017)) including i) tightening of the ligand binding site (LBS), ii) switching of the connector region, i.e. the P-I-F motif, into an active conformation, and iii) the G-protein binding site (GBS) becoming accessible to the cytoplasm. Nonetheless, many details have yet to be clarified. For example, LBS tightening is often provoked by agonist hydrogen bonding to polar residues in the extracellular end of H5 (H5_EC_) (often at the 5.46 position (Ballesteros/Weinstein nomenclature (Ballesteros & Weinstein, 1995))) at the same time as agonist electrostatic interaction with Asp3.32 (Latorraca et al., 2017; Vass et al., 2019). However, in a few cases LBS tightening has been observed to occur via H3, H6, or H7 movement in μ-opioid, muscarinic M2, and α-adrenergic receptors, respectively (Huang et al., 2015; Kruse et al., 2013; Yuan et al., 2020). Agonists incapable of equivalent polar interactions, outside of a charged amine anchored to Asp3.32, are rarer but exist and the mechanism presented above does thus not suit all agonists. To the best of our knowledge, only one study has observed evidence of an initial activation mechanism not involving polar interactions with H5 as part of the mechanism (Yuan et al., 2020).

**Figure 1.**
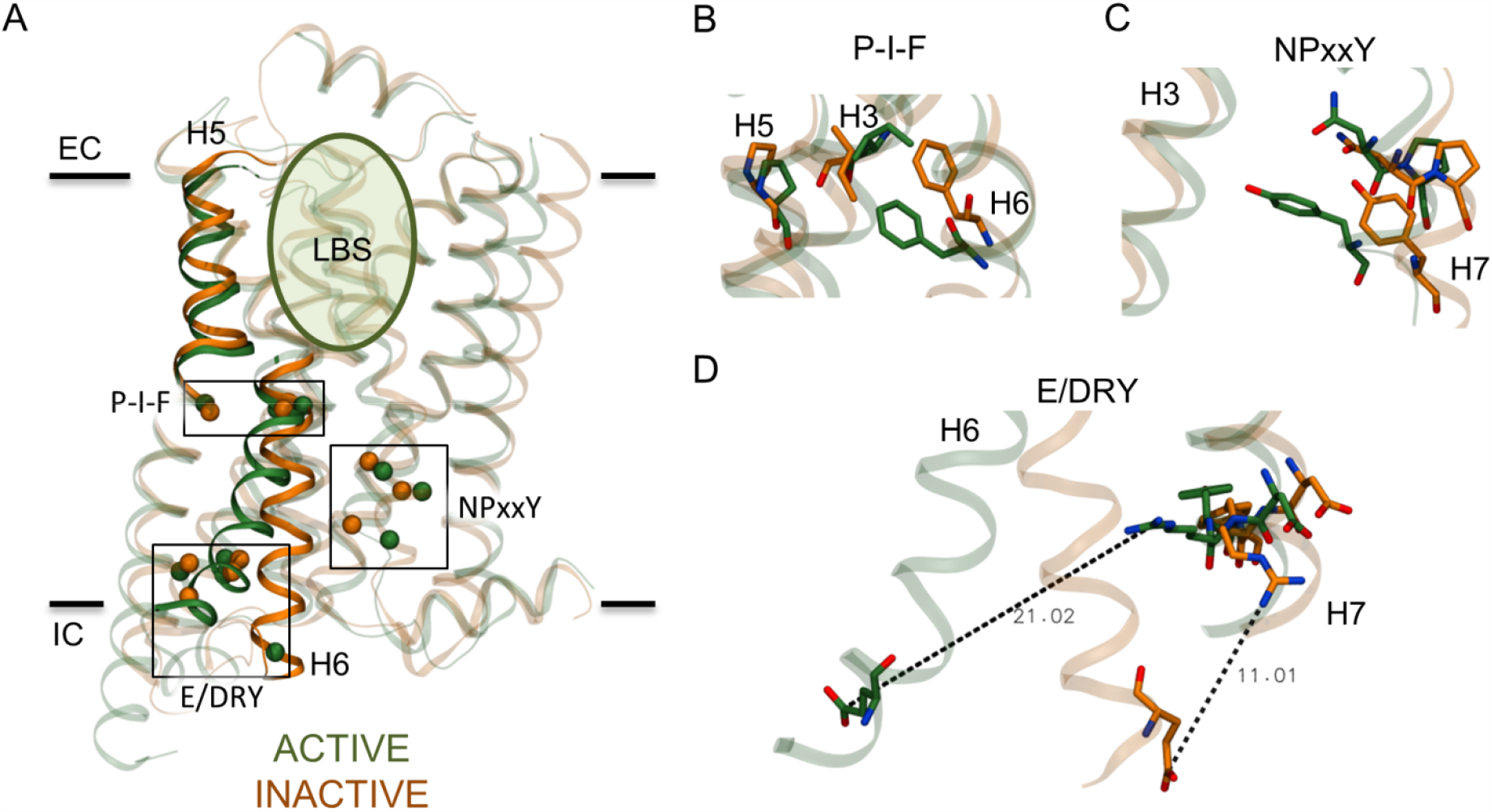
Overview of conformational changes in structural motifs between active (green) and inactive conformations (orange) of class A GPCRs. **A**) The LBS is indicated by a shaded circle in the protein structure, and the approximate location of the membrane is indicated by black lines. The EC and IC side of the membrane is noted. The H5_EC_ bulge and H6_IC_, which moves away from the protein core during activation, are shown in opaque ribbons. Important structural motifs, which undergo conformational changes upon activation, are indicated by boxes and opaque spheres at the Cα position of relevant residues. **B**) The connector region is centered around the P-I-F motif in helices H3, H5, and H6, and the active and inactive conformations are shown. The GBS is affected by changes in the **C**) NPxxY and **D**) E/DRY motifs. The Glu6.30-Arg3.50 distance, part of the E/DRY motif and known as the ionic lock, is indicated by a dashed line. In the inactive conformation, a salt bridge is often formed between the Glu6.30-Arg3.50; however, this interaction was disrupted in the crystallized receptor construct (orange). Experiments have revealed that two inactive conformations exist alongside each other; predominantly one where the ionic lock is locked, but also one where it is not (Dror et al., 2009; Manglik et al., 2015). The figure is based on two crystal structures of the β_2_ adrenergic receptor (PDB IDs: 2RH1 (Cherezov et al., 2007) and 3SN6 (Rasmussen et al., 2011)).

Vortioxetine (also known as LuAA21004, Brintellix, and Trintellix; Figure 2A) is a multimodal drug, meaning it interacts directly with multiple targets which complement each other in terms of efficacy and/or tolerability (Richelson, 2013), approved for the treatment of major depressive disorder. It has nanomolar affinity at its multiple targets and acts as a competitive inhibitor at the serotonin transporter (SERT); an antagonist at serotonergic 5-HT_3_ ion channel receptors as well as serotonergic 5-HT_1D_ and 5-HT_7_ GPCRs; a partial agonist at serotonergic 5-HT_1B_ receptors; and a full agonist at serotonergic 5-HT_1A_ receptors (Bang-Andersen et al., 2011; Sanchez, Asin, & Artigas, 2015). Serotonergic GPCRs belong to the aminergic cluster of class A GPCRs (Fredriksson, Lagerström, Lundin, & Schiöth, 2003) and couple to the Gα_i/o_ family of G-proteins to exert their downstream effects (Hannon & Hoyer, 2008). Serotonergic GPCRs include 12 receptors in total (Fredriksson et al., 2003) with sequence identities ranging from 25 to 62 % (Table S1); however, vortioxetine does not interact with the remaining serotonergic receptors to any physiologically relevant extent (Bang-Andersen et al., 2011). Vortioxetine’s comprehensive pharmacological profile has proven effective *in vivo* with an improved side effect profile compared to other antidepressant pharmacotherapies (Al-Sukhni, Maruschak, & McIntyre, 2015; Baldwin et al., 2016). However, the molecular mechanism underlying vortioxetine’s action has only been determined for SERT (Andersen et al., 2015) and the ionotropic 5-HT_3_ receptors (Ladefoged et al., 2018).

**Figure 2.**
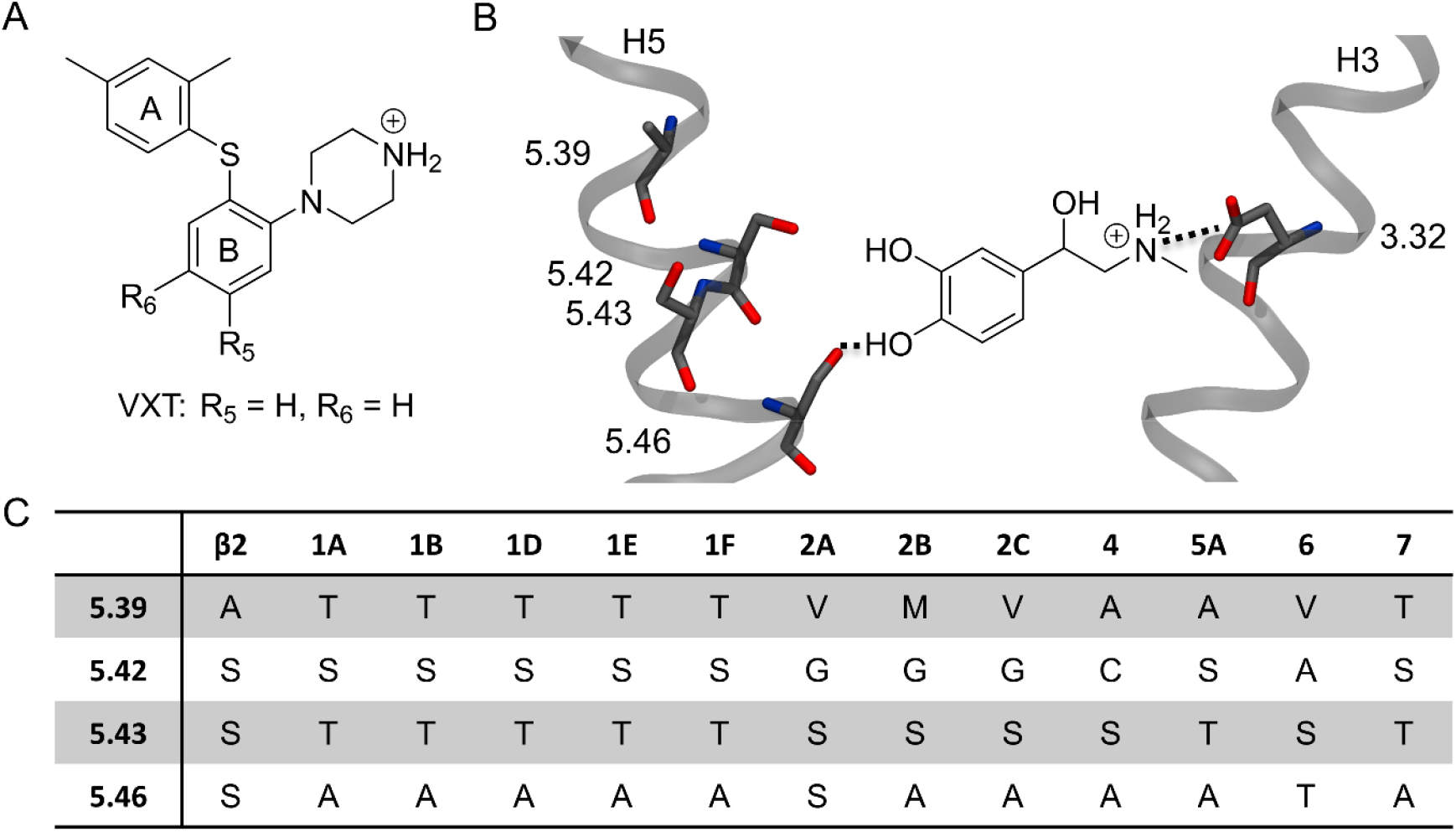
Receptor binding site and vortioxetine. **A)** the molecular structure of vortioxetine (VXT). The shown protonation state is that applied in the modeling. The two phenyl groups have been denoted phenyl A and B, and R_5_ and R_6_, used in the comparison to experimental structure-activity relationship data, are indicated on the structure. **B)** A schematic of the agonist/LBS interactions typically observed during activation in class A GPCRs as exemplified by the β2-adrenergic receptor (PDB-ID 3SN6). **C)** Amino acid conservation of residues 5.39, 5.42, 5.43, and 5.46 for the β2-adrenergic receptor and across the serotonergic GPCRs (Clustal Ω multiple sequence alignment).

Despite being a GPCR agonist, vortioxetine has no substituents capable of hydrogen bonding to H5 when the piperazine is anchored at Asp3.32, meaning vortioxetine necessarily activates the receptors using interactions distinct from the consensus activation mechanism. Additionally, in all serotonergic GPCRs, except for 5-HT_2A_ and 5-HT_6_, the residue at the 5.46 position is an alanine and is thus not able to participate in hydrogen bond interactions through the sidechain (Figure 2BC). Instead, residues at either one or two turns extracellular to residue 5.46, i.e. 5.43 and 5.39, respectively, are able to partake in hydrogen bond interactions in all serotonergic receptors. Investigation of vortioxetine binding and influence on the targeted GPCRs thus allows for exploration of i) how a small molecule can act as an agonist at one receptor while acting as an antagonist on a highly homologous receptor, and ii) how less polar agonists may activate GPCRs. Our findings considerably extend our current understanding of ligand recognition by all class A GPCRs and should improve future drug design efforts against these important targets.

## Results

The initial aim was to determine the most likely bioactive binding mode of vortioxetine in 5-HT_1A_, 5-HT_1B_, and 5-HT_7_ receptors in which vortioxetine acts as a full agonist, partial agonist, and antagonist, respectively (Bang-Andersen et al., 2011). The 5-HT_1D_ receptor was excluded from the series as vortioxetine binds to this target with lower affinity (Sanchez et al., 2015).

### Receptor modelling

In order to model vortioxetine binding in each receptor it is necessary to first obtain quality structures of each receptor in atomistic resolution. Optimally, each receptor structure should be in the conformational state stabilized by vortioxetine. Therefore, 5-HT_1A_ and 5-HT_1B_ should be in active conformations while 5-HT_7_ should be in an inactive conformation. At the time of modelling, only the 5-HT_1B_ structure had been solved experimentally. This structure is in a so-called active-like conformation (Wang et al., 2013) i.e. with an agonist bound in the LBS, but without a G-protein bound in the GBS. This structure was therefore used in the modelling of 5-HT_1B_, and also used as a template to model the 5-HT_1A_ receptor. A template for modelling the 5-HT_7_ receptor was searched for amongst all solved receptors from the aminergic cluster of class A GPCRs (Methods), and the most appropriate template was determined to be the dopamine D3 receptor in an inactive conformation.

The resulting models are shown in Figure 3 and Supporting Figure S1. It is clear that the 5-HT_1A_ and 5-HT_1B_ models are highly similar as expected based on the modelling approach, while the inactive 5-HT_7_ model is in another conformation. The conformation of H6_IC_ is similar in all three models due to the lack of bound G-protein in the active-like models. The LBS conformation, especially H5, H6, and H7, differs between active-like and inactive conformations (Figure 3) which will influence vortioxetine binding. Since modelling, the structure of a fully activated 5-HT_1B_ receptor has been solved (Garcia-Nafria, Nehme, Edwards, & Tate, 2018). A comparison of the LBS between structures reveals only minimal differences^1^, and the active-like models are thus appropriate for this study.

**Figure 3.**
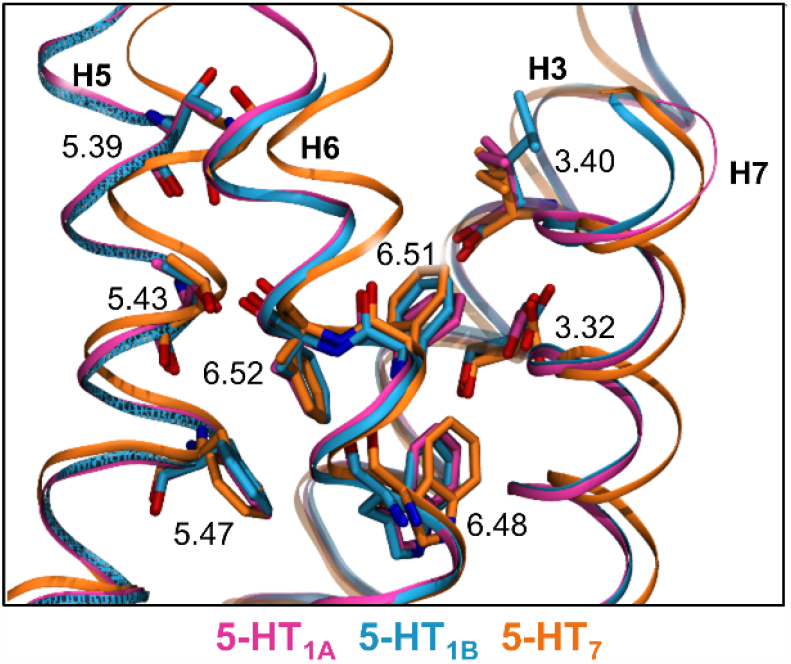
Comparison of the LBS of the three GPCR models; 5-HT_1A_ (pink), 5-HT_1B_ (blue), and 5-HT_7_ (orange). The 5-HT_1A_ and 5-HT_1B_ receptors are in active-like conformational states and the 5-HT_7_ receptor is in an inactive conformational state. The receptors are aligned on Cα atoms of H1-4.

### Determining the binding mode of vortioxetine in 5-HT_1A_, 5-HT_1B_, and 5-HT_7_ receptors

For each receptor, possible binding modes were generated *via* an induced fit docking calculation that incorporates both ligand and protein flexibility allowing for thorough binding mode exploration. The resulting poses (i.e. possible receptor/ligand complexes) were clustered based on the conformation and position of the ligand in the LBS. In order to select the bioactive binding mode from the several possible binding clusters it is necessary to implement additional methods. Two complementary methods were applied herein: i) the ligand stability was assessed for a representative pose from each binding cluster using short molecular dynamics simulations, and ii) the relative binding free energy between binding clusters were estimated using the molecular mechanics-Poisson Boltzmann and Surface Area (MM-PBSA) methodology. This approach has proven useful in selecting the bioactive binding mode of vortioxetine in the 5-TH_3_ ion channel receptor (Ladefoged et al., 2018) as well as in benchmarks covering a variation of protein targets (Liu & Kokubo, 2020).

The docking calculations resulted in several binding clusters for each receptor as shown in Figure 4 and summarized in Supporting Table S2. The cluster naming includes the receptor subtype in subscript to aid the reader. In all clusters, except for C4_7_, electrostatic interactions were detected between the charged amine of vortioxetine and the charged Asp3.32 in agreement with the consensus binding mechanism (Latorraca et al., 2017; Vass et al., 2019). Vortioxetine was able to bind in two overall orientations within the binding site in all three receptors. One orientation in which phenyl A of vortioxetine (Figure 2A) points toward the cytoplasm and is located furthest into the LBS (Figure 4ABC), and a flipped orientation in which phenyl B is pointing towards the cytoplasm and located furthest into the LBS (Figure 4DEF). It is intriguing that the ratio of the two overall orientations differ between the receptors, such that vortioxetine is primarily located with phenyl A pointing toward the intracellular side in the 5-HT_1A_ receptor in which vortioxetine acts as a full agonist, and phenyl A is pointing toward the extracellular side in the 5-HT_7_ receptor in which vortioxetine acts as an antagonist. Additionally, at the 5-HT_1B_ receptor, where vortioxetine acts as a partial agonist, the docking calculation proposes vortioxetine binding in both conformations close to the same extent. Albeit when accounting for the number of poses within each cluster for 5-HT_1B_, the orientation where phenyl A is pointing toward the intracellular side is the most prevalent orientation. It is also worth noting that in 5-HT_7_ vortioxetine generally occupies the space closer to the extracellular milieu in the LBS compared to vortioxetine docked into 5-HT_1A_ and 5-HT_1B_ regardless of orientation.

**Figure 4.**
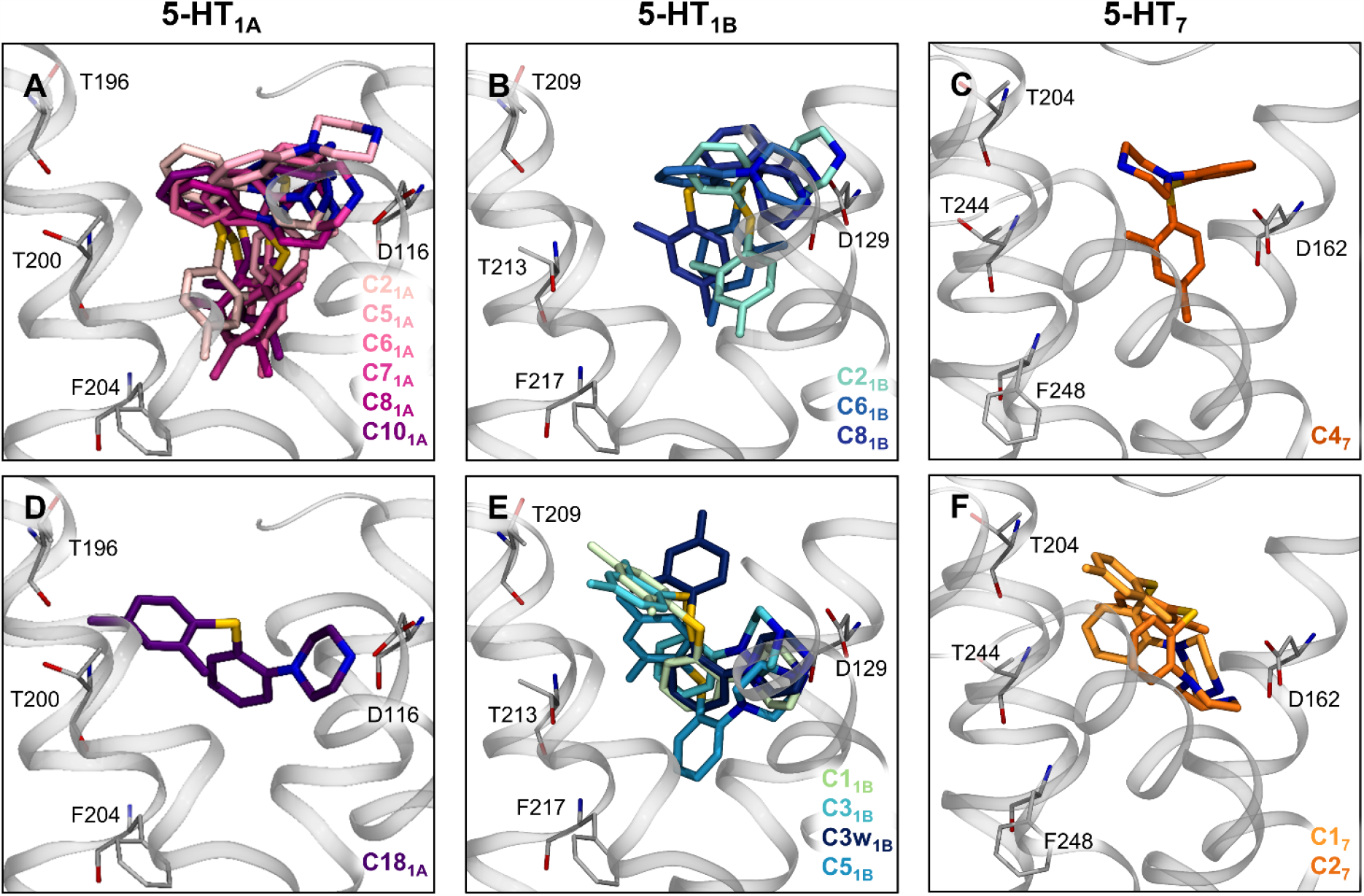
Binding modes of vortioxetine in the three receptors. Binding modes in the 5-HT_1A_ receptor are shown in pink hues, binding modes in the 5-HT_1B_ receptor are shown in blue hues, while binding modes in the 5-HT_7_ receptor are shown in orange hues. The view of the binding site is constant in all panels, and the changes in side chain orientations are due to the applied induced fit docking protocol. The binding clusters shown in each panel are listed in the bottom right corner of each panel using the same color as the corresponding vortioxetine molecule is shown in. **ABC)** A representative pose from each binding cluster of vortioxetine in which phenyl A is pointing towards the IC side of the protein for each receptor, 5-HT_1A_, 5-HT_1B_, and 5-HT_7_, respectively. **DEF)** Likewise, a representative pose from each binding cluster of vortioxetine in which phenyl A is oriented towards the EC side for the 5-HT_1A_, 5-HT_1B_, and 5-HT_7_ receptor, respectively.

The stability of each binding cluster was assessed using short, 50 ns MD simulations using three measures: i) the root-mean-squared deviation of vortioxetine (RMSD), ii) the fluctuation of vortioxetine along the membrane normal (i.e. the z-axis), and iii) the interatomic distance between vortioxetine’s charged amine and Asp3.32 (Figure 5). The most stable binding modes were then compared by their estimated relative binding free energy using MM-PBSA (see Methods and Supporting Table S3). The free energy was based on 100 snapshots evenly extracted from the initial 2 ns of the simulation time. By only using the first few nanoseconds the free energies can be used as a post-scoring method probing the energies of the docked pose, and not as an assessment of the MD-relaxed conformations of vortioxetine.

**Figure 5.**
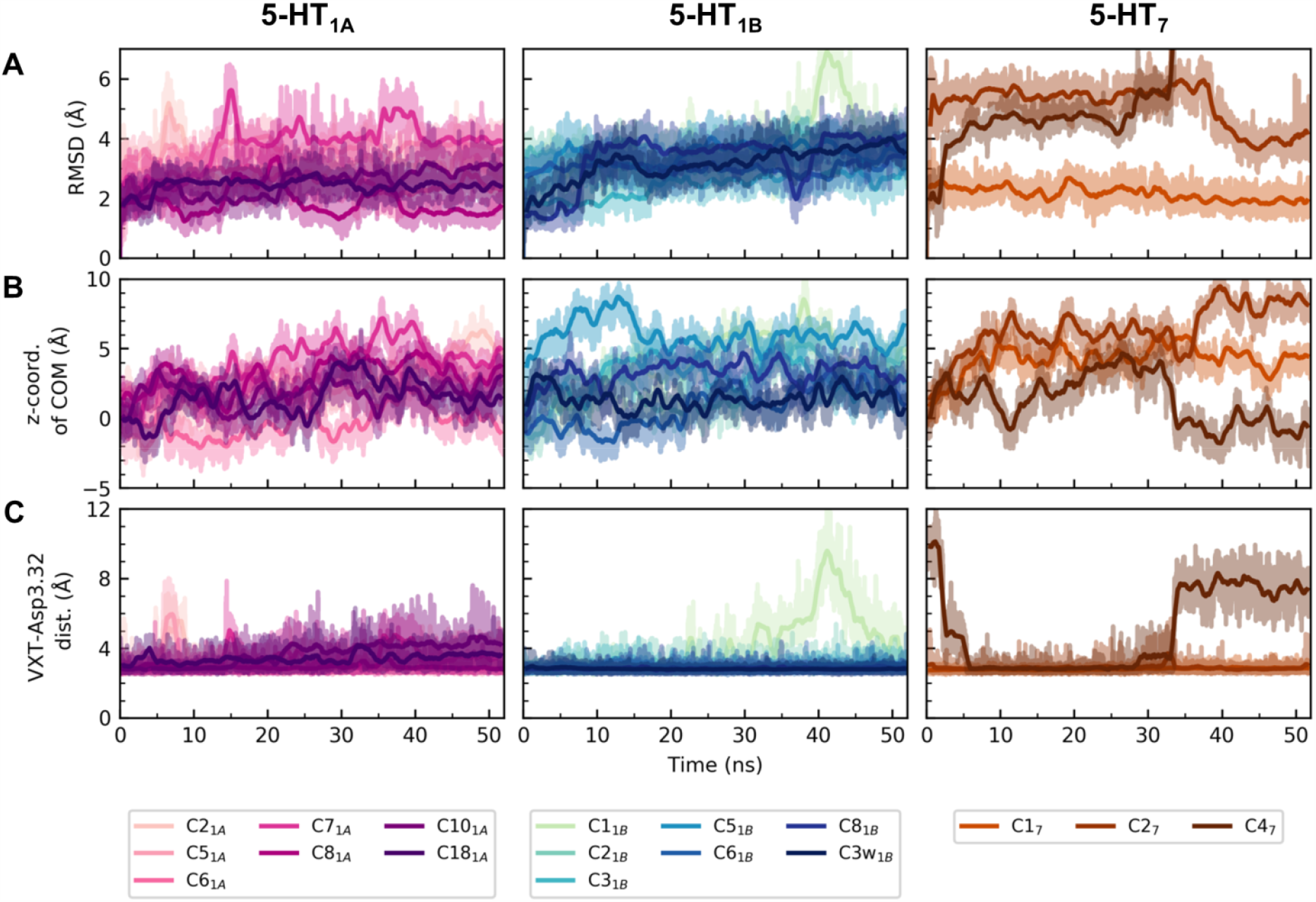
Stability measures of vortioxetine in each receptor. **A**) RMSD of vortioxetine (VXT) heavy atoms. **B**) The z-coordinate (membrane normal) of the center-of-mass (COM) of vortioxetine relative to the starting position. **C**) The distance between the charged amine in vortioxetine and the charged Asp3.32. This distance was calculated as the minimal heteroatom distance between amine and either carboxylate oxygen atom. The raw data is shown in transparent hues, while the running average is shown in opaque hues. The individual RMSD plots can be found in Supporting Figure S2, S3, and S4.

The MD simulations revealed that clusters identified for vortioxetine binding in 5-HT_1A_ and 5-HT_1B_ are more stable than 5-HT_7_. It was found that a given binding cluster had one of three fates: i) retaining the original binding mode; ii) evolving into an alternative cluster observed in the docking data; and iii) evolving into an entirely new binding mode (Supporting Figure S2, S3, and S4).

Using the above selection approach, clusters C6_1A_ and C8_1A_ are likely representatives of the bioactive binding mode of vortioxetine in 5-HT_1A_. It is possible that the bioactive conformation is an equilibrium between these two conformations. This idea is supported by the simulations, which show the only difference between the binding mode in C6_1A_ and C8_1A_ to be a 180° rotation of phenyl A by the end of the simulations (Supporting Figure S2CD). However, when taking the slight instability of C8_1A_ into account, C6_1A_ remains as the most likely bioactive binding mode. Upon closer inspection of the C6_1A_ simulation it was observed that H6_IC_ was pushed out in a manner similar to what occurs during GPCR activation (Latorraca et al., 2017). Signs of initial activation were not observed in any of the other simulations (Supporting Figure S5).

Some clusters were less stable in 5-HT_1B_ compared to vortioxetine binding to the 5-HT_1A_ receptor (Figure, Supporting Figure S3). Brief, partial unbinding was observed for C1_1B_ after approximately 40 ns, but vortioxetine returned to its initial binding mode before the end of the simulation. The analyses indicate C6_1B_ is the most likely bioactive binding mode.

In 5-HT_7_, only the binding mode in C1_7_ was observed to be stable (Figure, Supporting Figure S4). In C2_7_, vortioxetine moves even further toward the extracellular milieu, indicative of unbinding, and in C4_7_, vortioxetine moves in between H6_EC_ and H7_EC_ toward the membrane interior. The MM-PBSA calculations also show C1_7_ to be the most stable binding mode (Supporting Table S3). Thus based on the data at hand it is proposed that C1_7_ represents the bioactive binding mode of vortioxetine in the 5-HT_7_ receptor.

The binding cluster most likely to represent the bioactive binding mode of vortioxetine in each receptor is shown in Figure 6. In all three receptors, the charged amine in vortioxetine interacts electrostatically with Asp3.32. In 5-HT_1A_, phenyl A of vortioxetine is located at the very bottom of the LBS near Trp358_6.48_ and Phe204_5.47_ with the *ortho*-methyl group pointing towards H7, while phenyl B is located more extracellularly in the LBS lining residues Thr196_5.39_ and Thr200_5.43_. In 5-HT_1B_, vortioxetine is similarly located in the LBS compared to in the 5-HT_1A_ receptor, albeit approximately 1 Å higher up, with phenyl A interacting with Ile130_3.32_, Ile137_3.40_, and Phe331_6.52_. The main interaction partner of phenyl B is Phe330_6.51_. Lastly, in 5-HT_7_, the binding mode of vortioxetine is reversed such that phenyl A points to the extracellular side instead of the cytoplasm. Phenyl A is located close to H7 and is able to form cation-π interactions with Arg350_6.58_ and also interact hydrophobically with Leu232 from extracellular loop 2 (EL2), Phe343_6.51_, Leu346_6.54_, and Leu370_7.39_. Phenyl B is closer to H5, but interacts with Ile233_EL2_, Ile159_3.29_, Val163_3.33_, Phe343_6.51_, and Phe344_6.52_ and is additionally able to form π-π interactions with Phe343_6.51_.

**Figure 6.**
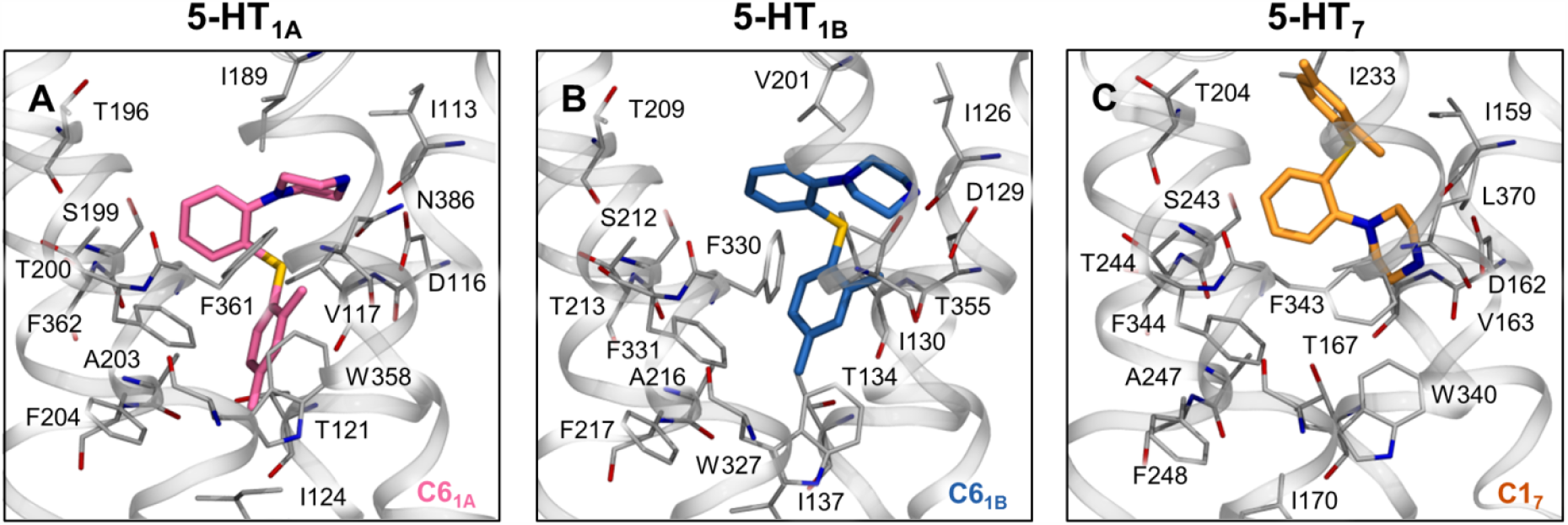
The proposed bioactive binding modes of vortioxetine in **A**) 5-HT_1A_, **B**) 5-HT_1B_, and **C**) 5-HT_7_ receptors. Residues which form the binding site are shown in gray tube, and H3, H5, H6, and H7 are shown as transparent ribbons.

Why vortioxetine occupies the binding site of 5-HT_7_ in an entirely different manner compared to 5-HT_1A_ and 5-HT_1B_ is intriguing and led us to delve deeper into an LBS comparison. A change in binding mode may be due to a i) change in amino acid sequence far from the LBS indirectly influencing the LBS conformation, and/or a ii) change of residue type within the LBS that directly perturb receptor/ligand interactions. Firstly, based on the homology models, H5_EC_, H6_EC_, and H7_EC_ are differently located in 5-HT_7_ compared to 5-HT_1A_ and 5-HT_1B_ (Figure 3), which influences the possible binding modes of vortioxetine. This could be an artefact of the homology modeling; however, the difference is also observed in the experimentally determined active and inactive conformations of 5-HT_1B_ (PDB ID: 6G79 and 5V54, respectively). Secondly, only five residues are not conserved in the LBS between the three receptors (3.28, 3.29, 3.33, 6.55, and 7.39, Supporting Table S4), and only one of these differ between 5-HT_7_ and both 5-HT_1A_ and 5-HT_1B_, namely residue 7.39 which is an asparagine in 5-HT_1A_, a threonine in 5-HT_1B_, and a leucine in 5-HT_7_. A hydrogen bond between the charged amine of vortioxetine and Asn7.39 was observed to be able to form simultaneously with the expected salt bridge between the same amine of vortioxetine and Asp3.32 in MD simulations of 5-HT_1A_/vortioxetine and 5-HT_1B_/vortioxetine complexes (Supporting Figure S6). It would therefore be of great interest to determine the functional effect of vortioxetine binding to mutant L7.39N and L7.39T 5-HT_7_ receptors.

### The influence of vortioxetine on receptor dynamics

Each of the bioactive binding modes identified above were further assessed using longer time-scale MD simulations with the aim of identifying how vortioxetine is capable of acting as either an agonist or an antagonist at the three receptors. Each receptor was simulated for 500 ns in three repeats (MD1-3), and evidence of receptor activation in the LBS, connector region, and GBS was monitored for each trajectory (Figure 7, Methods). According to large-scale MD simulations of adrenergic GPCRs (Dror et al., 2011; Kohlhoff et al., 2014), the events denoting receptor activation often occur independently in short periods of time. However, the fully activated conformational state arises when all events occur simultaneously and even then the GBS is only stably open when a G-protein is bound (Dror et al., 2011; Manglik et al., 2015).

**Figure 7.**
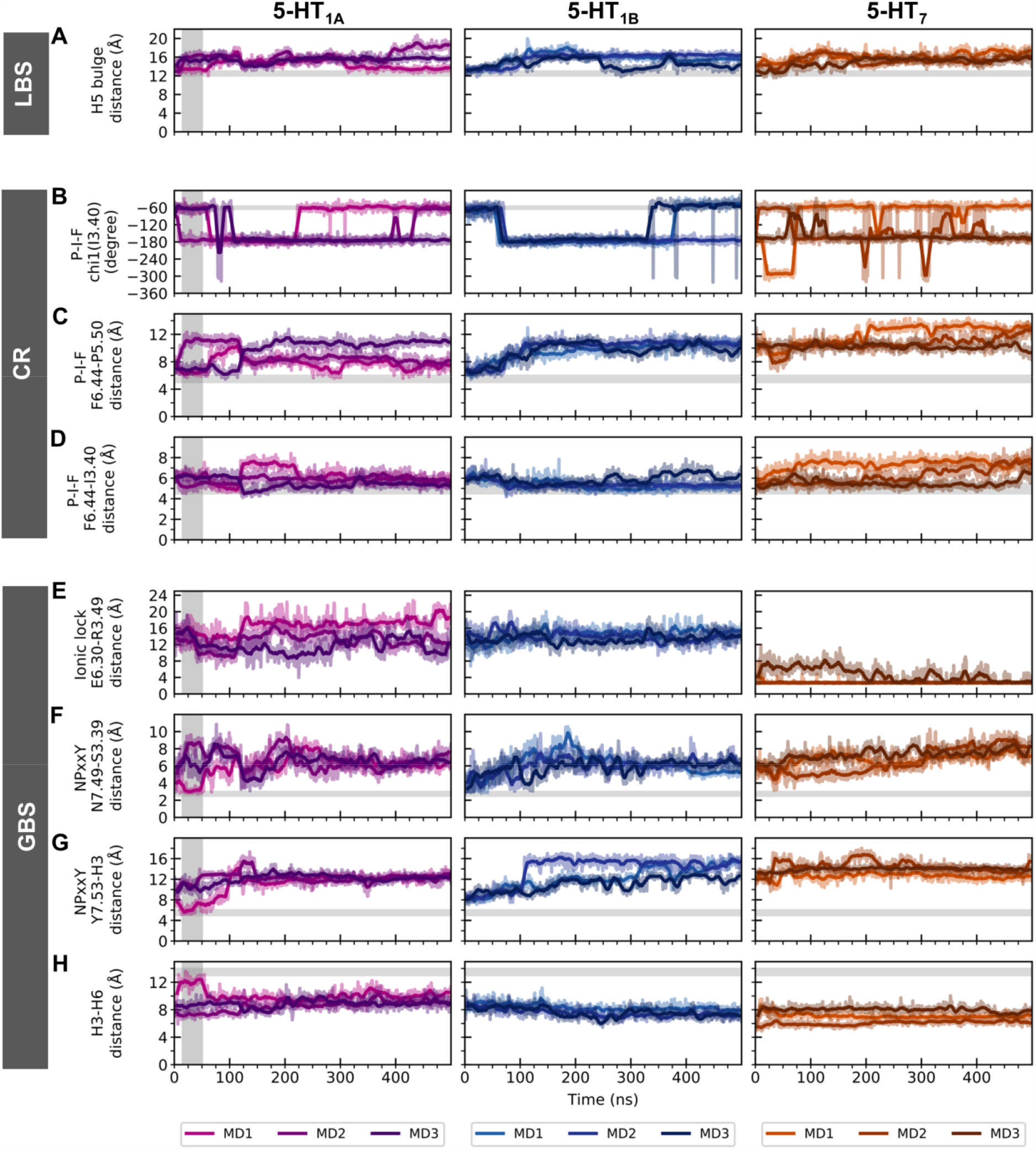
Measures of receptor activation. For 5-HT_1A_ (left, purple hues), 5-HT_1B_ (middle, blue hues), and 5-HT_7_ (right, orange hues), the time-progressed degree or distance changes are shown for **A**) the LBS, **B-D**) the connector region (CR), and **E-H**) the GBS. A vertical gray bar indicates the brief coordinated activation event by the 5-HT_1A_ receptor in the MD1 repeat. In each panel, the active-state value of each measure is indicated by a horizontal, gray line, except for the ionic lock panel, which could take on any value above 4 Å when in an active state. For each panel, the raw data is shown in transparent and the smoothed data is shown in opaque hues. Please see Methods for details.

During the simulations of all receptor/vortioxetine complexes, independent conformational changes to the monitored measures were observed as expected. However, all measures simultaneously switched to active conformations in the 5-HT_1A_ receptor during a brief ∼35 ns period in the beginning of MD1 (Figure 7A-H, vertical gray bar). The short time-span of the activation is in accord with the lack of a G-protein in the simulations, albeit shorter than what has previously been measured in experiments (Vilardaga, Bunemann, Krasel, Castro, & Lohse, 2003); however, the detection-limit of experiments is typically in the low ms-range. Thus, vortioxetine briefly activates the receptor in this simulation repeat. As 5-HT_1A_ is a constitutively active receptor (Berg & Clarke, 2018) the activation cannot be accredited to vortioxetine binding without further analysis. In fact, Yin et al. (2018) have performed MD simulations of 5-HT_1B_ in an inactive-like conformation using a model based on a similar chimeric crystallographic construct as was used herein, and observed that the model rapidly changed into a fully inactive conformation during simulations. The authors argued that the crystallized receptor was prevented from being fully inactive by the chimeric crystallization construct, but was crystallized in a so-called inactive-prone conformation, meaning that removal of the chimeric addition would lead to the rapid change into a fully inactive conformation during MD simulations (Yin et al., 2018). To assess if a conceptually similar active-prone conformation is inherent in the 5-HT_1A_ receptor model used herein, simulations of the *apo* 5-HT_1A_ model were performed in three repeats of 500 ns. In all repeats, 5-HT_1A_ was observed to retain its initial conformation, and no further activation or deactivation dynamics were observed (Supporting Figure S7). Collectively, this indicates that the brief activation observed in the 5-HT_1A_/vortioxetine MD1 simulation could be invoked by the presence of vortioxetine.

In 5-HT_7_, vortioxetine appears to stabilize an alternative inactive conformation compared to the conformation used in the docking study as H6_EC_ and H7_EC_ were observed to move away from the protein core during all repeat simulations (Figure 7A). The final conformation of 5-HT_7_ thus has an even wider ligand binding site compared to the initial receptor model. It is generally acknowledged that the inactive conformation of a GPCR consists of an ensemble of inactive conformations (Berg & Clarke, 2018), but this particular conformation has, to the best of our knowledge, not been observed in experimentally determined structures of antagonist/inverse agonist bound aminergic GPCRs^2^. The changes in conformation of H6_EC_ and H7_EC_ are thus not indicative of an unstable receptor, but may rather reflect different inactive conformations.

### Altered allosteric communication in 5-HT_1A_ during activation

In an effort to further investigate the activation mechanism of the 5-HT_1A_ receptor, the dynamical network methodology (Sethi, Eargle, Black, & Luthey-Schulten, 2009) was applied. This method assumes allosteric communication transfer occurs *via* non-bonded interactions between residues in a protein. The probability of information transfer between two residues is based on the (anti)correlation of movement of the residues in question (Methods). Each residue is mapped as a node and the inter-node communication is denoted an edge. The weight of an edge denotes the strength of communication between the nodes and is determined based on the degree of (anti)correlated motion between them.

The allosteric network can be grouped into communities of high internal communication. The communities assigned to 5-HT_1A_ in each repeat simulation primarily differ only in one region of the protein (Figure 8A). In MD1, in which the receptor briefly activates, H6_IC_ resides in the same community as the entirety of H5, while in MD2 and MD3, it does not. In MD2, the whole of H6 is in a community by itself, while in MD3, it is split in two; however, in neither of these two simulation repeats is H6 coupled to H5. This indicates that the movement of the intracellular end of H6 is tied to the movement of the intracellular end of H5 upon activation, but not otherwise. Furthermore, based on the node edges which link H5 to H6 and their assigned weights, this connection is strong (Figure 8A). Seven edges in MD1 are connecting H6_IC_ to H5_IC_, and they are significantly stronger than all other inter-helical edge weights within the receptor.

**Figure 8.**
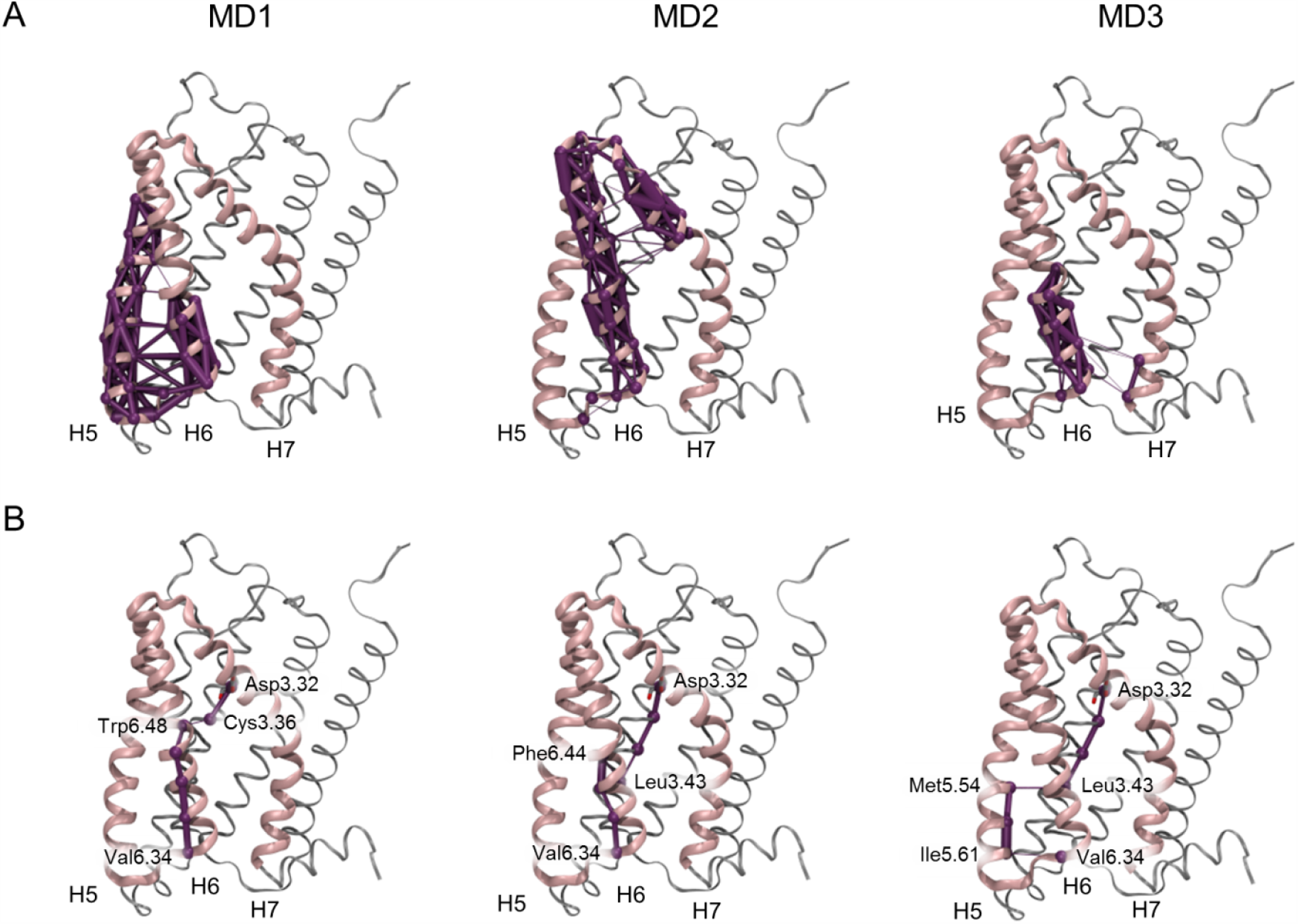
Dynamical network analysis of the 5-HT_1A_/vortioxetine complexes. **A)** the community which involves the intracellular end of H6 is shown for each simulation repeat, MD1-MD3. The edges which connect nodes within each community are shown as purple lines and their thickness represents their weights i.e. their strength of communication. **B)** the optimal path between Asp3.32, which anchors the bound agonist, and Val6.34, which is located at the most intracellular end of H6, is shown in purple for each repeat, and the residues bridging helices are named. The drawn nodes are weighted by the correlation strength between the two nodes such that thick lines denote highly (anti)correlated movement. H5, H6, and H7 are shown in pink ribbons, while the remainder of the protein is shown in gray ribbons.

The communication between LBS and GBS was assessed by determining the most optimal pathways of communication from Asp116_3.32_, which anchors the agonist to the LBS, to Val344_6.34_ located at the most intracellular end of H6 where the movement of H6_IC_ away from the protein core is most pronounced.

It is evident that the shortest path in MD1 is different compared to the paths determined for MD2 and MD3 (Figure 8B). The shortest path goes through the LBS (from Cys3.36 to Trp6.48) in MD1, where activation was observed, but not in the other repeats. Here the shortest path travels down H3 until below the LBS before jumping to H5 or H6. The paths thus indicate that the LBS is not directly involved in the communication pathway in the latter repeat simulations. Importantly, the altered allosteric communication pathway results in a shorter and stronger communication path between LBS and GBS during activation.

## Discussion

We applied docking, MD simulations, and free energy calculations to determine vortioxetine binding in 5-HT_1A_, 5-HT_1B_, and 5-HT_7_ receptors. In the first two receptors, where vortioxetine acts as an agonist, vortioxetine was found to occupy the lowest part of the LBS, while in the latter receptor, where vortioxetine is an antagonist, an alternative binding mode was observed. Extended MD simulations of the bioactive 5-HT_1A_/vortioxetine complex revealed an activation event invoked by vortioxetine.

The determined bioactive binding mode of vortioxetine in 5-HT_1A_ can be validated against pre-existing structure/activity relationship (SAR) and metabolite activity data (Bang-Andersen, Jørgensen, Bundgaard, Jensen, & Sanchéz, 2015; Bang-Andersen et al., 2011). A “methyl walk” performed on phenyl B of vortioxetine indicated that this phenyl is likely interacting closely with residues from the binding site (Bang-Andersen et al., 2011). Furthermore, the addition of a hydroxyl group to R_5_ or R_6_ (Figure 2A) on phenyl B abolishes binding affinity at 5-HT_1A_ (Bang-Andersen et al., 2015). In the determined bioactive binding mode, phenyl B is located close to H5; less than 4 Å away from Ser199_5.42_ and Thr200_5.43_ (shortest distance between heavy atoms). Therefore, even though the introduction of a hydrogen bond donor on phenyl B appears to be favorable considering its close proximity to polar residues, it appears that there is not enough space to accommodate it in this particular complex. More variation of the substituents on phenyl A is tolerated compared to phenyl B; however, the addition of more polar substituents affects binding affinity negatively. As the bottom of the binding site is an overall hydrophobic environment, the SAR data is in agreement with the determined binding mode. Surprisingly, however, vortioxetine metabolites with –CH_2_OH substituents on phenyl A instead of the methyl group are tolerated. It is possible that the CH_2_OH substituent placed in an *ortho*-position is able to reach Asp116_3.32_ in the determined binding mode; however, the affinity of vortioxetine with a *para*-substituted –CH_2_OH group cannot be explained by the determined binding mode. It is possible that water molecules might be bridging an interaction or that the addition of a larger, polar functional group alters the binding mode as has previously been observed for polar vortioxetine analogs in SERT (Andersen et al., 2015).

Unfortunately, no relevant experimental mutation data for the 5-HT_1B_ and 5-HT_7_ receptors was available to validate the binding modes.

The bioactive binding modes provide insight into how vortioxetine may act as an agonist at one receptor while acting as an antagonist on a highly homologous receptor. The determined binding modes of vortioxetine in 5-HT_1A_ and 5-HT_1B_ receptors are highly similar (Figure 6): the charged amine forms electrostatic interactions with Asp3.32, while both phenyl A and B line H5. Phenyl A is located furthest into the binding site where it can interact hydrophobically, and sometimes *via* π/π interactions, with Ile3.32, Ile3.40, Phe5.47, Trp6.48, Phe6.51, and Phe6.52. Overall, vortioxetine appears able to reach ∼1 Å deeper into the LBS in 5-TH_1A_ compared to 5-HT_1B_. In accordance with these observations, aromatic interactions with Trp6.48, Phe6.51, Phe6.52, and Tyr6.55 were detected in the experimentally solved α_2B_ adrenergic receptor complexed with an agonist unable to partake in hydrogen bonding with H5 (Yuan et al., 2020). In contrast, vortioxetine binds higher up the LBS in 5-HT_7_. Here phenyl A can form cation-π interactions with Arg6.58, while phenyl B is near EL2, H3_EC_, and H6_EC_. Based on our analyses, mutants V7.39N and V7.39T may be able to shift the function of vortioxetine from antagonist to agonist at the 5-HT_7_ receptor. According to Vass et al. (2019), ligand affinity is generally affected in the reverse 5-HT_1A_ N7.39V mutant and the conceptually similar 5-TH_1B_ T7.39N mutant; however, to the best of our knowledge, it has not been reported in the literature if the proposed mutants can alter the functional effect of a ligand.

The extended MD simulations revealed activation of the 5-HT_1A_ receptor. Recently, an experimentally solved 5-HT_1B_/agonist/Gα_o_-protein complex has been published (Garcia-Nafria et al., 2018), and the two active states were therefore compared. Despite being overall similar, several interesting differences were observed between the two active conformations. Firstly, the simulated outward movement of H6_IC_ was not as pronounced as detected in the 5-HT_1B_/agonist/Gα_o_-protein complex as well as other experimentally determined structures of GPCR/Gα_i/o_ complexes (Draper-Joyce et al., 2018; Garcia-Nafria et al., 2018; Kang et al., 2018; Kato et al., 2019; Koehl et al., 2018; Rasmussen et al., 2011). In these structures, an expansion of ∼8 Å is common between H3 and H6, while in MD1 herein an expansion of only ∼4 Å was observed (Figure 9A,C), indicating that the activation state we observe is not entirely comparable to G-protein bound structures. However, a study by Gregorio et al. (2017) monitored the H6_IC_ movement of β_2_-adrenergic receptors using single-molecule FRET analysis during the G-protein binding process. While they detected the expected 14 Å expansion in GPCR/agonist/Gα_s_ complex, they also detected two conformations of a GPCR/agonist complex; one of which has no shift of H6_IC_ compared to the inactive conformation, and one which has an up to 4 Å shift depending on the efficacy of the agonist. We propose that the active conformation detected herein for the 5-HT_1A_/vortioxetine complex is equivalent to the latter state, and thus, to the best of our knowledge, represents the first direct observation of this semi-active conformational state in atomistic detail. Secondly, during the activation event smaller changes to H7_IC_ and H8 were detected. In some experimentally determined structures H7_IC_ and H8 collectively move closer to the receptor core during activation (Draper-Joyce et al., 2018; Kang et al., 2018). In the simulations herein, partial movement of H7_IC_ and H8 was observed such that the part of H7 closest to the NPxxY motif had shifted, but H8 had not followed along (Figure 9B,C). We propose this to be an effect of the semi-active conformational state observed, and speculate that the complete movement would occur upon full activation following G-protein binding.

**Figure 9.**
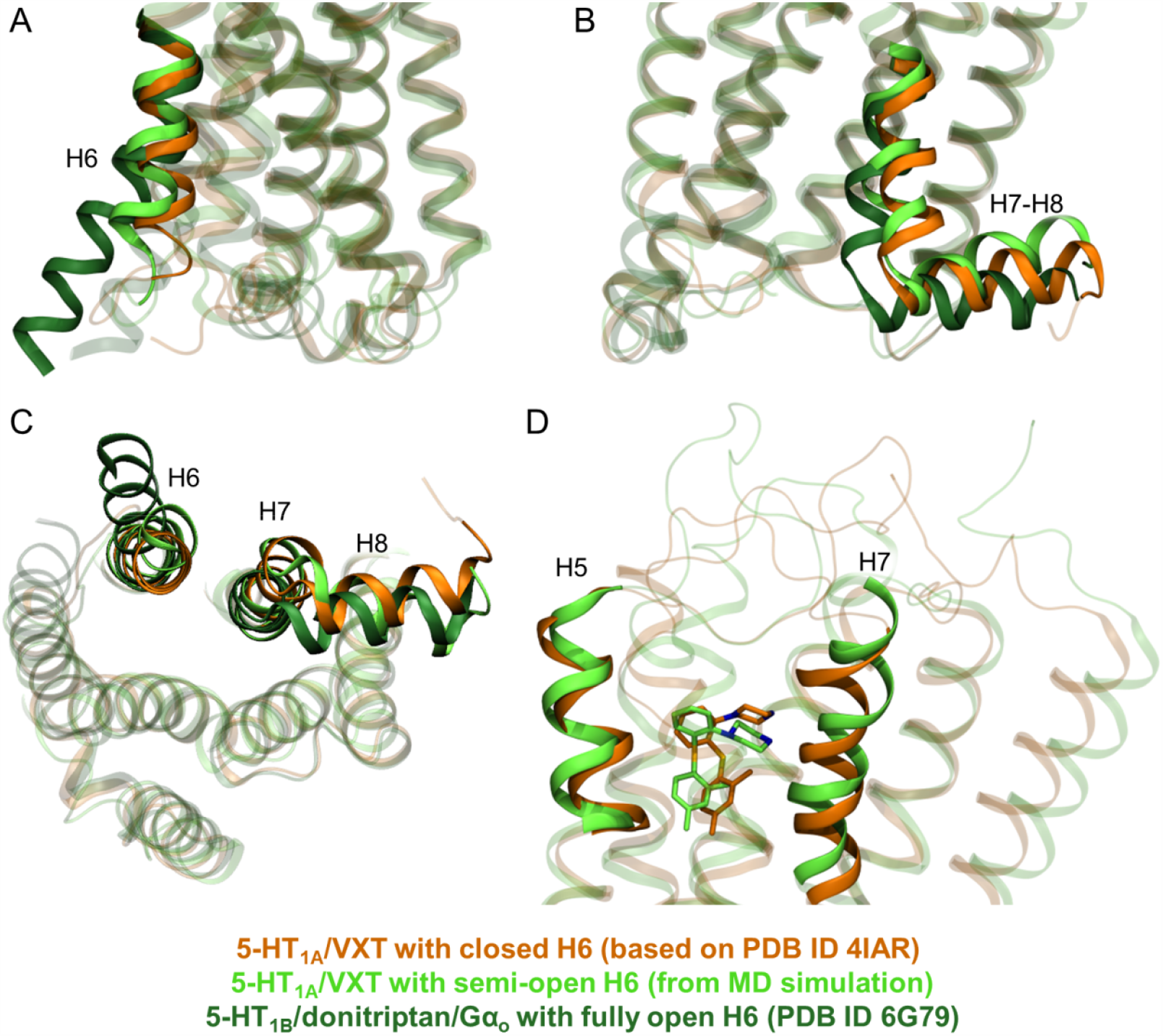
Comparison of the most active 5-HT_1A_ conformation observed in the MD1 simulation (light green) to the original 5-HT_1A_ homology model (orange) as well as a fully active 5-HT_1B_ receptor (dark green, based on PDB ID: 6G79 (Garcia-Nafria et al., 2018)). Comparisons of the conformation of **A**) H6_IC_ as well as **B**) H7_IC_ and H8 are shown as seen from within the membrane. **C**) An intracellular view of H6_IC_, H7_IC_, and H8 are shown. **D**) A sideview comparison of the local conformation of H5_EC_ and H7_EC_ are shown as well as the conformation of vortioxetine in each overall conformational state.

Differences in LBS conformation compared to the consensus activation mechanism were also detected. Generally, slight bending of H5_EC_ occurs during activation such that the H5_EC_-H7_EC_ distance becomes shorter. In the 5-HT_1A_/vortioxetine complex, where the intramolecular interactions are hydrophobic and aromatic rather than polar, no such bending was observed (Figure 9D). In fact, H5_EC_ was observed to move further away from the receptor core. Nonetheless, the calculated H5_EC_-H7_EC_ distance was shorter during activation, and closer inspection of H7_EC_ revealed a shift toward the receptor core. Intriguingly, this observation is in full accord with the α_2B_ adrenergic receptor complexed with an agonist lacking hydrogen bonding ability in the ligand core (Yuan et al., 2020), and thus hints toward a general initial step of the activation mechanism by less polar agonists. It is further intriguing that the 7.39 residue hypothesized to be important for vortioxetine function is located in H7_EC_ important for LBS tightening during agonist activation of 5-HT_1A_ in the MD simulations.

The dynamical network analysis revealed several residues important for allosteric communication between the LBS and the most intracellular end of H6 (Figure 8B). Due to the abundance of mutagenesis data for GPCRs (Vass et al., 2019) it is possible to validate these results by comparison of the hotspot residues with receptor responses to mutation of said residues. The allosteric communication pathway determined for MD1, in which activation was detected, jumps from Cys3.36 to Trp6.48. According to experiments, Cys3.36 can interact directly with Trp6.48 in the inactive state (Nichols & Nichols, 2008; Trzaskowski et al., 2012), and C3.36A mutation results in a loss of agonist effect by serotonin (Nichols & Nichols, 2008). Additionally, the conformation of the Trp6.48 side chain has been implicated in the activation mechanism of class A GPCRs (Trzaskowski et al., 2012). The experimental data thus fully support the role of Cys3.36 and Trp6.48 during activation. An alternative pathway was observed in the remaining simulations. Here the pathway jumps from Leu3.43 to Phe6.44 or Met5.54. Mutation of Phe6.44 into leucine or tyrosine has been observed to increase and decrease agonist potency in adrenergic receptors, respectively (Greasley, Fanelli, Rossier, Abuin, & Cotecchia, 2002). Phe6.44 is located at the midpoint of H6 and its nearby environment is altered upon activation when the intracellular end of H6 moves away from the protein core. The mutation’s effect on agonist potency could therefore also be explained by de- or increasing this residue’s interaction energy with nearby residues whereby making the receptor more or less prone to activation, which would be in accord with the calculated communication pathways.

We set out to determine vortioxetine binding at 5-HT_1A_, 5-HT_1B_, and 5-HT_7_ receptors with the aim of exploring i) how a small molecule can act as an agonist in one receptor and as an antagonist in a highly homologous receptor, and ii) how less polar agonists may activate GPCRs. Vortioxetine was determined to bind in opposite orientations and with differing interaction patterns in receptors in which it acts as an agonist *versus* receptors in which it acts as an antagonist, and our analyses indicate residue 7.39 to be important for the functional effect of vortioxetine. We have provided additional support of an activation mechanism utilized by less polar agonists proposed by (Yuan et al., 2020) in which aromatic interactions with residues at the bottom of the LBS substitute polar interactions in H5. Finally, the extended MD simulations revealed vortioxetine-invoked receptor activation of 5-HT_1A_, and further inspection of the event uncovered the finding of an as-of-yet unobserved step in the activation pathway: a receptor/agonist complex with a semi-open H6_IC_ ready for G-protein binding and further activation.

## Methods

### Protein preparation

The 5-HT_1B_ receptor model was based on the crystal structure of 5-HT_1B_ with co-crystallized ergotamine (PDB ID: 4IAR (Wang et al., 2013)). Two structural water molecules were found in the additionally published structure, 4IAQ, which were included in the 5-HT_1B_ model. The thermostable mutant W138L was mutated back to its native form, and missing atoms in residues were built using Prime 3.7 from the Schrödinger Suite 2014 (Schrödinger LLC, New York, NY). EL2 (Lys191-Ser197) and EL3 (Cys340-Cys344) were modeled ab initio using the extended serial loop sampling protocol in Prime 3.7 (Zhu, Pincus, Zhao, & Friesner, 2006). An implicit membrane and the VSGB water model was applied, and 10 loop conformations were produced for each loop. The lowest energy conformation was selected for each loop. Intracellular loop 3 (IL3) was too long to model and the loop ends (Arg238 and Ala306) were instead covalently linked due to their close proximity. The three loops, EL2, EL3, and IL3 were then minimized using OPLS-2005 (Kaminski, Friesner, Tirado-Rives, & Jorgensen, 2001). The resulting model was prepared using the Protein Preparation Wizard (Sastry, Adzhigirey, Day, Annabhimoju, & Sherman, 2013), PROPKA (Olsson, Søndergaard, Rostkowski, & Jensen, 2011), and prior knowledge of the protonation state of key residues during activation (Dror et al., 2011; Vogel et al., 2008). Asp146_3.49_ and Glu309_6.30_ were modeled as neutral, and His81_2.36_ and His347_7.31_ were modeled as the ε-tautomer. A disulfide bridge was included between Cys122_3.25_ and Cys199_EL2_. All other residues were modeled in their default states. The resulting model was validated by its ability to correctly predict the known binding mode of ergotamine using IFD calculations, and produce models of 5-HT and LSD binding that are consistent with available data (Wang et al., 2013).

The 5-HT_1A_ receptor (uniprot ID: P08908) was modeled based on the 5-HT_1B_ receptor model described above. The sequences were aligned using the structural alignment methodology with GPCR restraints in MOE 2013 (Chemical Computing Group ULC, Canada) (Supporting Figure S8). 100 backbone models were created using MOE 2013, and a further 10 models including side chains were created for each backbone model, resulting in 1000 models total. The optimal model was determined by its ability to correctly dock a library of 952 5-HT_1A_ agonists (Gatica & Cavasotto, 2012) and correctly exclude 37,128 decoys obtained from the ZINC database (Irwin, Sterling, Mysinger, Bolstad, & Coleman, 2012) during virtual screening using Glide 6.4 with SP-scoring from the Schrödinger Suite 2014 (Schrödinger LLC, New York, NY). All ligands were prepared using LigPrep from the Schrödinger Suite using pH = 7 and the OPLS-2005 force field (Kaminski et al., 2001). The enrichment factor (EF) (Kirchmair, Markt,

Distinto, Wolber, & Langer, 2008) was then calculated for each model, and models with EF > 4.5 at 1% of the dataset were further evaluated. The resulting eight models were compared structurally, and were found to be highly similar (RMSD < 2 Å for heavy atoms within both backbone and side chains). The eight models were evaluated based on Ramachandran plot violations, total number of agonists retrieved during screening, and the area under the receiver operator characteristic (ROC) curve (Kirchmair et al., 2008). The best model was prepared as described for 5-HT_1B_ above. Asp133_3.49_ and Glu340_6.30_ were modeled as neutral, His126_3.42_ was modeled as the ε-tautomer, and a disulfide bridge was modeled between Cys109_3.25_ and Cys187_EL2_. All other residues were modeled in their default states. The model was additionally validated by its ability to dock 5-HT, ergotamine, and LSD in a manner consistent with available data (Wang et al., 2013).

The 5-HT_7_ receptor (uniprot ID: P34969) was modeled based on the dopamine D3 receptor (PDB ID: 3PBL (Chien et al., 2010)). This template was chosen because it is crystallized in an inactive conformation, shares 35% sequence identity, and best reproduces the location of proline residues in the helical segments compared to other available aminergic GPCRs at the time of modeling (uniprot IDs: P07550, P07700, P35462, P08483, P08172, and P35367, i.e. adrenergic β_1_ and β_2_, dopaminergic D3, muscarinic M2 and M3, and histidinic H1 receptors), which has been shown to be important for constructing most accurate GPCR models (Hall, Roberts, & Vaidehi, 2009). The initial alignments were constructed using clustal Ω 2.1 (Sievers et al., 2011), while the final alignment was constructed using AlignMe (Stamm, Staritzbichler, Khafizov, & Forrest, 2013), including a secondary structure prediction and manually adjusted (Supporting Figure S8). 100 models were built using MODELLER 9.16 (Sali & Blundell, 1993) while enforcing a disulfide bridge between Cys155_3.25_ and Cys231_EL2_. The optimal model was selected based on DOPE and molpdf scores and Ramachandran plot violations. The resulting model was lacking the extracellular end of H1 which was then manually constructed by enforcing α-helical restraints in Maestro 9.9 from the Schrödinger Suite 2014 (Schrödinger LLC, New York, NY). The short IL3 and EL3 loops were optimized using the serial loop sampling protocol (Jacobson et al., 2004) and the OPLS-2005 force field (Kaminski et al., 2001) in Prime 3.7. The resulting model was prepared as described for 5-HT_1B_ above. All amino acids were modeled as their default states. The final model was validated by its ability to predict a model of 5-HT binding consistent with available data (Wang et al., 2013) based on the IFD protocol.

### Ligand preparation

The chemical structure of ergotamine, LSD, and 5-HT were built in Maestro 9.9 from the Schrödinger Suite 2014 (Schrödinger LLC, New York, NY), while vortioxetine was prepared as described in (Andersen et al., 2015). All structures were minimized using a conjugate gradient method and the OPLS 2.1 force field (Shivakumar et al., 2010), and the protonation states were assessed using Epik (Greenwood, Calkins, Sullivan, & Shelley, 2010; Shelley et al., 2007), All ligands were modeled as having a formal charge of +1. The ligands were then subjected to a conformational search algorithm using a mixed torsional/low mode sampling algorithm, which combines Monte Carlo torsional sampling and exploration of low-frequency eigenvectors to sample conformations, and the OPLS 2.1 force field (Shivakumar et al., 2010) in MacroModel 11.4. The lowest energy conformation of each ligand was used in docking calculations.

### Docking

All docking calculations were performed using the induced fit docking protocol (Sherman, Day, Jacobson, Friesner, & Farid, 2006) employing Glide 6.4 and Prime 3.7 in the Schrödinger Suite 2014. In the initial docking all vdW interactions were scaled to 50%, the SP level was applied, and a maximum of 200 poses were allowed. The binding site was defined as the geometric centroid of Asp3.32, Phe6.51, and Ala5.46 in calculations using the 5-HT_1A_ and 5-HT_7_ receptor models, while the co-crystallized granisetron was used for 5-HT_1B_ receptor docking calculations. In the optimization step, residues within 5 Å of vortioxetine were subjected to side chain optimization. The final docking step was performed in XP, and a maximum of 100 poses with associated energies within 30 kcal/mol of the lowest energy pose were reported in the results. The resulting poses were clustered based on their in-place conformation using the conformer cluster script available in Maestro 9.9 from the Schrödinger Suite 2014 (Schrödinger LLC, New York, NY). All clusters with fewer than two poses were considered outliers of the calculations, except when the average XP Gscore of the cluster was lower than the average XP Gscores of any large cluster.

### MD simulations

Simulation systems were built and equilibrated using a combined coarse grained (CG)/all-atom (AA) approach before production runs were performed in AA resolution. Each AA receptor was aligned on the OPM (Lomize, Pogozheva, Joo, Mosberg, & Lomize, 2012) structure of 5-HT_1B_ (PDB ID: 4IAR) and converted into CG resolution. A CG POPC membrane was built around the protein in the xy plane using Insane (Wassenaar, Ingolfsson, Bockmann, Tieleman, & Marrink, 2015) and Martinize tools available from the Marrink group’s homepage (http://cgmartini.nl/index.php/tools2/proteins-and-bilayers). The system was then solvated and ionized with NaCl to a concentration of 0.2 M and equilibrated while using position restraints on the protein according to step 1 and 2 in Table 1. The system was then converted into its atomistic equivalent using the Backward tool (Wassenaar, Pluhackova, Bockmann, Marrink, & Tieleman, 2014). In order to ensure that the starting point of the AA simulations was exactly as intended during protein preparation, the backmapped protein was replaced by the original receptor/vortioxetine complex. The AA system was then minimized using a conjugate gradient algorithm and further equilibrated in AA resolution before production runs, as outlined in step 3 to 5 in Table 1.

**Table 1.** Simulation protocol for receptor systems. The duration of the production runs were 50 ns for the short simulations of all receptor/vortioxetine clusters, and 500 ns for the most likely receptor/vortioxetine binding modes.

|  | Equilibration |  |  |  |  | Production run |
| --- | --- | --- | --- | --- | --- | --- |
| Sequential step | 1 | 2 | 3 | 4 | 5 | 6 |
| Resolution | CG | CG | AA | AA | AA | AA |
| Ensemble | NVT | NPT | NVT | NPT | NPT | NPT |
| Duration | 0.5 ns | 5 ns | 0.5 ns | 1 ns | 2 ns | 50 ns/500 ns |
| Timestep | 10 fs | 10 fs | 1 fs | 2 fs | 2 fs | 2 fs |
| Position restraints | Protein | Protein | Protein +ligand | Protein +ligand | None | None |
| Thermostat | Velocity rescale | Velocity rescale | Velocity rescale | Nose-Hoover | Nose-Hoover | Nose-Hoover |
| Barostat | - | Berendsen | - | Parrinello-Rahman | Parrinello-Rahman | Parrinello-Rahman |

During analysis, the trajectories obtained in step 5 and 6 (Table 1) were combined for all short simulations of possible binding modes, resulting in 52 ns simulations, as the binding mode of vortioxetine was observed to change within the initial 2 ns of unrestrained simulation time. The most likely binding mode of vortioxetine in each receptor was chosen for extension. For the most likely binding mode, the equilibrated system resulting from step 4 (Table 1) was extended for 500 ns in three repeats using different initial velocities.

All simulations were performed in Gromacs 5.0.2 (Abraham et al., 2015) using periodic boundary conditions. For the CG simulations, the MARTINI 2.2 force field (de Jong et al., 2013) was used. The vdW interactions were treated using cut-offs at 11 Å and the potential-shift-Verlet modifier, while electrostatic interactions were treated using the reaction-field method cut-off at 11 Å and a dielectric constant of 0 (= infinite) beyond the cut-off. The neighbor list was maintained using Verlet buffer lists. Temperature was kept at 310 K and pressure at 1 bar. The AA resolution simulations were performed using the CHARMM36 force field (Best et al., 2012; Klauda et al., 2010; MacKerell et al., 1998), the TIPS3P water model (Durell, Brooks, & Bennaim, 1994), and CHARMM-compatible vortioxetine parameters as described in (Ladefoged et al., 2018). The vdW interactions were treated by cut-offs at 12 Å and a force-switch modifier after 10 Å, while electrostatic interactions were treated using PME. The neighbor list was maintained using Verlet buffer lists, and bonds linking hydrogen atoms to heavy atoms were restrained using LINCS (Hess, 2008). The temperature was maintained at 310 K using a coupling constant of 1, and pressure was maintained at 1 bar using a coupling constant of 4, a compressibility factor of 4.5 x10-5 and semi-isotropic coupling to xy and z dimensions separately.

### Measures of activation

The conformational states of the receptors during MD simulations were assessed by the conformation of multiple conserved motifs and helix segments. The tightening of the ligand binding site was calculated as the distance between Cα atoms of residues 5.46 and 7.42 in agreement with (Cherezov et al., 2007; Rasmussen et al., 2011). Changes in conformation of residues in the P-I-F motif which form the connector region were also monitored. Based on GPCR structures available in the PDB at the time of analysis (PDB IDs: 3PBL, 4IB4, 4IAR, 4IAQ, 5D5A, 5D5B, 4QKX, 4LDE, 4LDL, 4LDO, 4GBR, 3SN6, 3P0G, 3PDS, 3NY8, 3NY9, 3NYA, 3D4S, 2R4R, 2R4S, and 2RH1), the χ1 angle of Ile3.40 (calculated for atoms N-Cα-Cβ-Cγ1) was observed to be gauche+ in the active conformation and anti in the inactive conformation. Additionally, the location of Phe6.44 was observed to shift relative to Ile3.40 and Pro5.50, such that in the active conformation the phenylalanine was closer to Pro5.50 compared to in the inactive conformation. The Cα distances between both Pro5.50 and Phe6.44 as well as Ile3.40 and Phe6.44 were therefore calculated. In the GBS multiple motifs were assessed. The distance between the polar side chain of Glu6.30 and Arg3.50 was calculated as the minimum distance between either the oxygen or nitrogen atom in the side chain of glutamate and arginine, respectively. The conformational change of the NPxxY motif was assessed in two ways: *i)* the shortest distance between polar atoms of the side chain of Asn7.49 and Ser3.39, as a hydrogen bond between these two residues have been observed to stabilize an active receptor conformation (Kohlhoff et al., 2014), and *ii)* the proximity of Tyr7.53 to the H3-H5-H6 bundle (Erlandson, McMahon, & Kruse, 2018). The proximity was calculated as the distance between Cα atoms of Tyr7.53 and residue 3.44, which was chosen due to its placement being similar to Tyr7.53 regarding the membrane normal, and its placement in the conformationally stable H3. Lastly, the movement of the intracellular end of H6 away from the rest of the protein was calculated as the distance between Cα atoms of residues 3.50 and 6.35 (Cherezov et al., 2007; Rasmussen et al., 2011).

### MM-PBSA calculation

Binding free energy calculations were performed for each receptor/vortioxetine cluster using GMXPBSA 2.1 (Paissoni, Spiliotopoulos, Musco, & Spitaleri, 2014, 2015) based on 100 frames evenly extracted from the first 2 ns of the 52 ns production run MD simulations. The snapshots were thus extracted directly after ligand restraints had been removed such that the calculated binding free energies could be used as a post-docking scoring method, but not as an assessment of the binding mode each complex converged towards during the short MD simulations. The 1-MD approach, in which all calculations are based on a single MD simulation instead of several, was applied and the molecular system was stripped of lipid, ions, and water before the calculations. The software APBS (Baker, Sept, Joseph, Holst, & McCammon, 2001) was used to solve the Poisson-Boltzmann equation and calculate the nonpolar contribution to the solvation energy, while Gromacs 5.0.2 (Abraham et al., 2015) and the CHARMM36 force field (Best et al., 2012; MacKerell et al., 1998) was used to calculate the vdW and electrostatic energy contribution. The Poisson-Boltzmann equation was solved using a non-linear approximation and “SDH” boundary conditions. A grid spacing of 1, a temperature of 310 K, an ion concentration of 0.2 M, and a protein dielectric constant of 2 was applied. The “multitrj” option was applied to ensure use of the exact same grid definitions across the different receptor/vortioxetine complexes.

### Dynamical network analysis

The probability of information transfer i.e. allostery between two residues is based on the (anti)correlation of movement of the residues in question. For each repeat of the 5-HT_1A_/vortioxetine simulations, each Cα atom was mapped as a node, and non-neighboring nodes were connected by so-called edges. Edges were assigned a weight based on the magnitude of the calculated correlation in motion between the two nodes. Based on the weights, the nodes were divided into communities representing substructures in the protein which are more densely interconnected compared to the rest of the protein (Sethi et al., 2009). (Sub)optimal pathways of communication are calculated as the shortest distance between the two residues in terms of edge weights (Sethi et al., 2009).

The dynamical networks were visualized using NetworkView in VMD 1.9.3 (Humphrey, Dalke, & Schulten, 1996). The correlation of atom motions were calculated using CARMA (Glykos, 2006), and communities and suboptimal paths were calculated using scripts obtained from the Schulten group webpage (http://faculty.scs.illinois.edu/schulten/Software2.0.html#4) (gncommunities and subopt, respectively). Only protein heavy atoms were included in the calculation, and nodes were assigned to Cα atoms. The correlated motion of atoms within the same residue and the nearest neighbor were excluded from the analysis. The suboptimal paths linking Asp3.32 to Val6.34 were calculated using an edge length offset of 20 i.e. the length of the longest suboptimal path must be less than 20 longer than the optimal path.

## Acknowledgements

We would like to thank Peter Mathias Nemec Nielsen for performing IFD calculations on the 5-HT_1B_ structure, and Anne Laustsen for providing the 5-HT_1A_ homology model. The research was funded by the Carlsberg Foundation (CF15-0572), the Lundbeck Foundation (R126-A12453) and the Independent Research Fund Denmark | Medical Sciences (DFF – 4004-00309). Computational resources were provided from the Novo Nordisk Foundation (NF18OC0032608). All computations were made possible at the Grendel-cluster through the Centre for Scientific Computing, Aarhus (http://www.cscaa.dk/grendel/hardware/).

## Footnotes

1 Root-mean-squared deviation < 0.5 Å for Cα atoms within intracellular ends of H1-H7.

2 Visual comparison of H6_EC_-H7_EC_ to the following PDB structures (aligned on H1-4 Cα atoms): 5V54, 6A94, 6DRX, 6BQH, 4BVN, 2YCZ, 2YCW, 5×7D, 3NY8, 3NY9, 3NYA, 6PRZ, 2VT4, 2YCX, 2YCY, 5A8E, 6DS0, 6DRZ, 6A93, 6PS0, 6PS1, 6PS4, 6PS5, 2RH1, 2R4R, 2R4S, 6CM4, 3PBL, 6IQL, 3RZE, 5CXV, 5YC8, 3UON, 5ZHP, 4U14, 4U15, 4U16, 4DAJ, 5DSG, and 6OL9.

## References

Abraham, M. J., Murtola, T., Schulz, R., Páll, S., Smith, J. C., Hess, B., & Lindahl, E. (2015). GROMACS: high performance molecular simulations through multi-level parallelism from laptops to supercomputers. SoftwareX, 1-2, 19-25. doi:https://doi.org/10.1016/j.softx.2015.06.001

Al-Sukhni, M., Maruschak, N. A., & McIntyre, R. S. (2015). Vortioxetine: a review of efficacy, safety and tolerability with a focus on cognitive symptoms in major depressive disorder. Expert Opinion on Drug Safety, 14(8), 1291–1304. doi:10.1517/14740338.2015.1046836

Andersen, J., Ladefoged, L. K., Wang, D. Y., Kristensen, T. N. B., Bang-Andersen, B., Kristensen, A. S., Schiøtt, B., & Strømgaard, K. (2015). Binding of the multimodal antidepressant drug vortioxetine to the human serotonin transporter. ACS Chemical Neuroscience, 6(11), 1892–1900. doi:10.1021/acschemneuro.5b00225

Baker, N. A., Sept, D., Joseph, S., Holst, M. J., & McCammon, J. A. (2001). Electrostatics of nanosystems: application to microtubules and the ribosome. Proceedings of the National Academy of Sciences of the United States of America, 98(18), 10037–10041. doi:10.1073/pnas.181342398

Baldwin, D. S., Chrones, L., Florea, I., Nielsen, R., Nomikos, G. G., Palo, W., & Reines, E. (2016). The safety and tolerability of vortioxetine: analysis of data from randomized placebo-controlled trials and open-label extension studies. Journal of Psychopharmacology, 30(3), 242–252. doi:10.1177/0269881116628440

Ballesteros, J. A., & Weinstein, H. (1995). [19] Integrated methods for the construction of three-dimensional models and computational probing of structure-function relations in G protein-coupled receptors (Vol. 25): Academic Press.

Bang-Andersen, B., Jørgensen, M., Bundgaard, C., Jensen, C. G., & Sanchéz, C. (2015). Case history: brintellix® (vortioxetine, LU AA21004), an antidepressant with multimodal activity. Medicinal Chemistry Reviews, 50, 433–445.

Bang-Andersen, B., Ruhland, T., Jørgensen, M., Smith, G., Frederiksen, K., Jensen, K. G., Zhong, H. L., Nielsen, S. M., Hogg, S., Mørk, A., & Stensbøl, T. B. (2011). Discovery of 1-[2-(2,4-dimethylphenylsulfanyl)phenyl]piperazine (Lu AA21004): a novel multimodal compound for the treatment of major depressive disorder. Journal of Medicinal Chemistry, 54(9), 3206–3221. doi:10.1021/jm101459g

Berg, K. A., & Clarke, W. P. (2018). Making sense of pharmacology: inverse agonism and functional selectivity. International Journal of Neuropsychopharmacology, 21(10), 962–977. doi:10.1093/ijnp/pyy071

Best, R. B., Zhu, X., Shim, J., Lopes, P. E., Mittal, J., Feig, M., & Mackerell, A. D. Jr., (2012). Optimization of the additive CHARMM all-atom protein force field targeting improved sampling of the backbone phi, psi and side-chain chi(1) and chi(2) dihedral angles. Journal of Chemical Theory and Computation, 8(9), 3257–3273. doi:10.1021/ct300400x

Cherezov, V., Rosenbaum, D. M., Hanson, M. A., Rasmussen, S. G., Thian, F. S., Kobilka, T. S., Choi, H. J., Kuhn, P., Weis, W. I., Kobilka, B. K., & Stevens, R. C. (2007). High-resolution crystal structure of an engineered human beta2-adrenergic G protein-coupled receptor. Science, 318(5854), 1258–1265. doi:10.1126/science.1150577

Chien, E. Y. T., Liu, W., Zhao, Q. A., Katritch, V., Han, G. W., Hanson, M. A., Shi, L., Newman, A. H., Javitch, J. A., Cherezov, V., & Stevens, R. C. (2010). Structure of the human dopamine D3 receptor in complex with a D2/D3 selective antagonist. Science, 330(6007), 1091–1095. doi:10.1126/science.1197410

de Jong, D. H., Singh, G., Bennett, W. F., Arnarez, C., Wassenaar, T. A., Schafer, L. V., Periole, X., Tieleman, D. P., & Marrink, S. J. (2013). Improved parameters for the MARTINI coarse-grained protein force field. Journal of Chemical Theory and Computation, 9(1), 687–697. doi:10.1021/ct300646g

Draper-Joyce, C. J., Khoshouei, M., Thal, D. M., Liang, Y. L., Nguyen, A. T. N., Furness, S. G. B., Venugopal, H., Baltos, J. A., Plitzko, J. M., Danev, R., Baumeister, W., May, L. T., Wootten, D., Sexton, P. M., Glukhova, A., & Christopoulos, A. (2018). Structure of the adenosine-bound human adenosine A1 receptor-Gi complex. Nature, 558(7711), 559–563. doi:10.1038/s41586-018-0236-6

Dror, R. O., Arlow, D. H., Borhani, D. W., Jensen, M. O., Piana, S., & Shaw, D. E. (2009). Identification of two distinct inactive conformations of the beta(2)-adrenergic receptor reconciles structural and biochemical observations. Proceedings of the National Academy of Sciences of the United States of America, 106(12), 4689–4694. doi:10.1073/pnas.0811065106

Dror, R. O., Arlow, D. H., Maragakis, P., Mildorf, T. J., Pan, A. C., Xu, H. F., Borhani, D. W., & Shaw, D. E. (2011). Activation mechanism of the beta2-adrenergic receptor. Proceedings of the National Academy of Sciences of the United States of America, 108(46), 18684–18689. doi:10.1073/pnas.1110499108

Durell, S. R., Brooks, B. R., & Bennaim, A. (1994). Solvent-induced forces between 2 hydrophilic groups. Journal of Physical Chemistry, 98(8), 2198–2202. doi:DOI 10.1021/j100059a038

Erlandson, S. C., McMahon, C., & Kruse, A. C. (2018). Structural basis for G protein-coupled receptor signaling. Annu Rev Biophys. doi:10.1146/annurev-biophys-070317-032931

Fredriksson, R., Lagerström, M. C., Lundin, L. G., & Schiöth, H. B. (2003). The G-protein-coupled receptors in the human genome form five main families. Phylogenetic analysis, paralogon groups, and fingerprints. Molecular Pharmacology, 63(6), 1256–1272. doi:DOI 10.1124/mol.63.6.1256

Garcia-Nafria, J., Nehme, R., Edwards, P. C., & Tate, C. G. (2018). Cryo-EM structure of the serotonin 5-HT1B receptor coupled to heterotrimeric Go. Nature, 558(7711), 620–623. doi:10.1038/s41586-018-0241-9

Gatica, E. A., & Cavasotto, C. N. (2012). Ligand and decoy sets for docking to G protein-coupled receptors. Journal of Chemical Information and Modeling, 52(1), 1–6. doi:10.1021/ci200412p

Glykos, N. M. (2006). Carma: a molecular dynamics analysis program. Journal of Computational Chemistry, 27(14), 1765–1768. doi:10.1002/jcc.20482

Greasley, P. J., Fanelli, F., Rossier, O., Abuin, L., & Cotecchia, S. (2002). Mutagenesis and modelling of the alpha1b-adrenergic receptor highlight the role of the helix 3/helix 6 interface in receptor activation. Molecular Pharmacology, 61(5), 1025-1032. Retrieved from https://www.ncbi.nlm.nih.gov/pubmed/11961120

Greenwood, J. R., Calkins, D., Sullivan, A. P., & Shelley, J. C. (2010). Towards the comprehensive, rapid, and accurate prediction of the favorable tautomeric states of drug-like molecules in aqueous solution. Journal of Computer-Aided Molecular Design, 24(6-7), 591–604. doi:10.1007/s10822-010-9349-1

Gregorio, G. G., Masureel, M., Hilger, D., Terry, D. S., Juette, M., Zhao, H., Zhou, Z., Perez-Aguilar, J. M., Hauge, M., Mathiasen, S., Javitch, J. A., Weinstein, H., Kobilka, B. K., & Blanchard, S. C. (2017). Single-molecule analysis of ligand efficacy in beta(2)AR-G-protein activation. Nature, 547(7661), 68-+. doi:10.1038/nature22354

Hall, S. E., Roberts, K., & Vaidehi, N. (2009). Position of helical kinks in membrane protein crystal structures and the accuracy of computational prediction. Journal of Molecular Graphics and Modelling, 27(8), 944–950. doi:10.1016/j.jmgm.2009.02.004

Hannon, J., & Hoyer, D. (2008). Molecular biology of 5-HT receptors. Behavioural Brain Research, 195(1), 198–213. doi:10.1016/j.bbr.2008.03.020

Hess, B. (2008). P-LINCS: a parallel linear constraint solver for molecular simulation. Journal of Chemical Theory and Computation, 4(1), 116–122. doi:10.1021/ct700200b

Huang, W. J., Manglik, A., Venkatakrishnan, A. J., Laeremans, T., Feinberg, E. N., Sanborn, L., Kato, H. E., Livingston, K. E., Thorsen, T. S., Kling, R. C., Granier, S., Gmeiner, P., Husbands, S. M., Traynor, J. R., Weis, W. I., Steyaert, J., Dror, R. O., & Kobilka, K. (2015). Structural insights into mu-opioid receptor activation. Nature, 524(7565), 315-+. doi:10.1038/nature14886

Humphrey, W., Dalke, A., & Schulten, K. (1996). VMD: Visual molecular dynamics. Journal of Molecular Graphics and Modelling, 14(1), 33–38. doi:DOI 10.1016/0263-7855(96)00018-5

Irwin, J. J., Sterling, T., Mysinger, M. M., Bolstad, E. S., & Coleman, R. G. (2012). ZINC: a free tool to discover chemistry for biology. Journal of Chemical Information and Modeling, 52(7), 1757–1768. doi:10.1021/ci3001277

Jacobson, M. P., Pincus, D. L., Rapp, C. S., Day, T. J. F., Honig, B., Shaw, D. E., & Friesner, R. A. (2004). A hierarchical approach to all-atom protein loop prediction. Proteins-Structure Function and Bioinformatics, 55(2), 351–367. doi:10.1002/prot.10613

Kaminski, G. A., Friesner, R. A., Tirado-Rives, J., & Jorgensen, W. L. (2001). Evaluation and reparametrization of the OPLS-AA force field for proteins via comparison with accurate quantum chemical calculations on peptides. Journal of Physical Chemistry B, 105(28), 6474–6487. doi:10.1021/jp003919d

Kang, Y., Kuybeda, O., de Waal, P. W., Mukherjee, S., Van Eps, N., Dutka, P., Zhou, X. E., Bartesaghi, A., Erramilli, S., Morizumi, T., Gu, X., Yin, Y., Liu, P., Jiang, Y., Meng, X., Zhao, G., Melcher, K., Ernst, O. P., Kossiakoff, A. A., Subramaniam, S., & Xu, H.E. (2018). Cryo-EM structure of human rhodopsin bound to an inhibitory G protein. Nature, 558(7711), 553–558. doi:10.1038/s41586-018-0215-y

Kato, H. E., Zhang, Y., Hu, H., Suomivuori, C. M., Kadji, F. M. N., Aoki, J., Krishna Kumar, K., Fonseca, R., Hilger, D., Huang, W., Latorraca, N. R., Inoue, A., Dror, R. O., Kobilka, B. K., & Skiniotis, G. (2019). Conformational transitions of a neurotensin receptor 1-Gi1 complex. Nature, 572(7767), 80–85. doi:10.1038/s41586-019-1337-6

Kirchmair, J., Markt, P., Distinto, S., Wolber, G., & Langer, T. (2008). Evaluation of the performance of 3D virtual screening protocols: RMSD comparisons, enrichment assessments, and decoy selection - what can we learn from earlier mistakes? Journal of Computer-Aided Molecular Design, 22(3-4), 213–228. doi:10.1007/s10822-007-9163-6

Klauda, J. B., Venable, R. M., Freites, J. A., O’Connor, J. W., Tobias, D. J., Mondragon-Ramirez, C., Vorobyov, I., MacKerell, A. D. Jr.,, & Pastor, R. W. (2010). Update of the CHARMM all-atom additive force field for lipids: validation on six lipid types. J Phys Chem B, 114(23), 7830–7843. doi:10.1021/jp101759q

Koehl, A., Hu, H., Maeda, S., Zhang, Y., Qu, Q., Paggi, J. M., Latorraca, N. R., Hilger, D., Dawson, R., Matile, H., Schertler, G. F. X., Granier, S., Weis, W. I., Dror, R. O., Manglik, A., Skiniotis, G., & Kobilka, B. K. (2018). Structure of the micro-opioid receptor-Gi protein complex. Nature, 558(7711), 547–552. doi:10.1038/s41586-018-0219-7

Kohlhoff, K. J., Shukla, D., Lawrenz, M., Bowman, G. R., Konerding, D. E., Belov, D., Altman, R. B., & Pande, V. S. (2014). Cloud-based simulations on Google Exacycle reveal ligand modulation of GPCR activation pathways. Nature Chemistry, 6(1), 15–21. doi:10.1038/Nchem.1821

Kruse, A. C., Ring, A. M., Manglik, A., Hu, J. X., Hu, K., Eitel, K., Hubner, H., Pardon, E., Valant, C., Sexton, P. M., Christopoulos, A., Felder, C. C., Gmeiner, P., Steyaert, J., Weis, W. I., Garcia, K. C., Wess, J., & Kobilka, B. K. (2013). Activation and allosteric modulation of a muscarinic acetylcholine receptor. Nature, 504(7478), 101-+. doi:10.1038/nature12735

Ladefoged, L. K., Munro, L., Pedersen, A. J., Lummis, S. C. R., Bang-Andersen, B., Balle, T., Schiøtt, B., & Kristensen, A. S. (2018). Modeling and mutational analysis of the binding mode for the multimodal antidepressant drug vortioxetine to the human 5-HT3A receptor. Molecular Pharmacology, 94(6), 1421–1434. doi:10.1124/mol.118.113530

Latorraca, N. R., Venkatakrishnan, A. J., & Dror, R. O. (2017). GPCR Dynamics: Structures in Motion. Chemical Reviews, 117(1), 139–155. doi:10.1021/acs.chemrev.6b00177

Liu, K., & Kokubo, H. (2020). Prediction of ligand binding mode among multiple cross-docking poses by molecular dynamics simulations. Journal of Computer-Aided Molecular Design. doi:10.1007/s10822-020-00340-y

Lomize, M. A., Pogozheva, I. D., Joo, H., Mosberg, H. I., & Lomize, A. L. (2012). OPM database and PPM web server: resources for positioning of proteins in membranes. Nucleic Acids Research, 40(D1), D370–D376. doi:10.1093/nar/gkr703

MacKerell, A. D., Bashford, D., Bellott, M., Dunbrack, R. L., Evanseck, J. D., Field, M. J., Fischer, S., Gao, J., Guo, H., Ha, S., Joseph-McCarthy, D., Kuchnir, L., Kuczera, K., Lau, F. T. K., Mattos, C., Michnick, S., Ngo, T., Nguyen, D. T., Prodhom, B., Reiher, W. E., Roux, B., Schlenkrich, M., Smith, J. C., Stote, R., Straub, J., Watanabe, M., Wiorkiewicz-Kuczera, J., Yin, D., & Karplus, M. (1998). All-atom empirical potential for molecular modeling and dynamics studies of proteins. Journal of Physical Chemistry B, 102(18), 3586–3616. doi:DOI 10.1021/jp973084f

Manglik, A., Kim, T. H., Masureel, M., Altenbach, C., Yang, Z. Y., Hilger, D., Lerch, M. T., Kobilka, T. S., Thian, F. S., Hubbell, W. L., Prosser, R. S., & Kobilka, B. K. (2015). Structural Insights into the Dynamic Process of beta(2)-Adrenergic Receptor Signaling. Cell, 161(5), 1101–1111. doi:10.1016/j.cell.2015.04.043

Nichols, D. E., & Nichols, C. D. (2008). Serotonin receptors. Chem Rev, 108(5), 1614–1641. doi:10.1021/cr078224o

Olsson, M. H. M., Søndergaard, C. R., Rostkowski, M., & Jensen, J. H. (2011). PROPKA3: consistent treatment of internal and surface residues in empirical pKa predictions. Journal of Chemical Theory and Computation, 7(2), 525–537. doi:10.1021/ct100578z

Paissoni, C., Spiliotopoulos, D., Musco, G., & Spitaleri, A. (2014). GMXPBSA 2.0: A GROMACS tool to perform MM/PBSA and computational alanine scanning. Computer Physics Communications, 185(11), 2920–2929. doi:10.1016/j.cpc.2014.06.019

Paissoni, C., Spiliotopoulos, D., Musco, G., & Spitaleri, A. (2015). GMXPBSA 2.1: a GROMACS tool to perform MM/PBSA and computational alanine scanning. Computer Physics Communications, 186, 105–107. doi:https://doi.org/10.1016/j.cpc.2014.09.010

Rasmussen, S. G., DeVree, B. T., Zou, Y., Kruse, A. C., Chung, K. Y., Kobilka, T. S., Thian, F. S., Chae, P. S., Pardon, E., Calinski, D., Mathiesen, J. M., Shah, S. T., Lyons, J. A., Caffrey, M., Gellman, S. H., Steyaert, J., Skiniotis, G., Weis, W. I., Sunahara, R. K., & Kobilka, B. K. (2011). Crystal structure of the beta2 adrenergic receptor-Gs protein complex. Nature, 477(7366), 549–555. doi:10.1038/nature10361

Richelson, E. (2013). Multi-modality: a new approach for the treatment of major depressive disorder. International Journal of Neuropsychopharmacology, 16(6), 1433–1442. doi:10.1017/S1461145712001605

Sali, A., & Blundell, T. L. (1993). Comparative protein modelling by satisfaction of spatial restraints. Journal of Molecular Biology, 234(3), 779–815. doi:10.1006/jmbi.1993.1626

Sanchez, C., Asin, K. E., & Artigas, F. (2015). Vortioxetine, a novel antidepressant with multimodal activity: review of preclinical and clinical data. Pharmacol Ther, 145, 43–57. doi:10.1016/j.pharmthera.2014.07.001

Santos, R., Ursu, O., Gaulton, A., Bento, A. P., Donadi, R. S., Bologa, C. G., Karlsson, A., Al-Lazikani, B., Hersey, A., Oprea, T. I., & Overington, J. P. (2017). A comprehensive map of molecular drug targets. Nature Reviews Drug Discovery, 16(1), 19–34. doi:10.1038/nrd.2016.230

Sastry, G. M., Adzhigirey, M., Day, T., Annabhimoju, R., & Sherman, W. (2013). Protein and ligand preparation: parameters, protocols, and influence on virtual screening enrichments. Journal of Computer-Aided Molecular Design, 27(3), 221–234. doi:10.1007/s10822-013-9644-8

Sethi, A., Eargle, J., Black, A. A., & Luthey-Schulten, Z. (2009). Dynamical networks in tRNA: protein complexes. Proceedings of the National Academy of Sciences of the United States of America, 106(16), 6620–6625. doi:10.1073/pnas.0810961106

Shelley, J. C., Cholleti, A., Frye, L. L., Greenwood, J. R., Timlin, M. R., & Uchimaya, M. (2007). Epik: a software program for pKa prediction and protonation state generation for drug-like molecules. Journal of Computer-Aided Molecular Design, 21(12), 681–691. doi:10.1007/s10822-007-9133-z

Sherman, W., Day, T., Jacobson, M. P., Friesner, R. A., & Farid, R. (2006). Novel procedure for modeling ligand/receptor induced fit effects. Journal of Medicinal Chemistry, 49(2), 534–553. doi:10.1021/jm050540c

Shivakumar, D., Williams, J., Wu, Y. J., Damm, W., Shelley, J., & Sherman, W. (2010). Prediction of absolute solvation free energies using molecular dynamics free energy perturbation and the OPLS force field. Journal of Chemical Theory and Computation, 6(5), 1509–1519. doi:10.1021/ct900587b

Sievers, F., Wilm, A., Dineen, D., Gibson, T. J., Karplus, K., Li, W. Z., Lopez, R., McWilliam, H., Remmert, M., Soding, J., Thompson, J. D., & Higgins, D. G. (2011). Fast, scalable generation of high-quality protein multiple sequence alignments using Clustal Omega. Molecular Systems Biology, 7. doi:ARTN 539 10.1038/msb.2011.75

Stamm, M., Staritzbichler, R., Khafizov, K., & Forrest, L. R. (2013). Alignment of helical membrane protein sequences using AlignMe. PloS One, 8(3). doi:ARTN e57731 10.1371/journal.pone.0057731

Trzaskowski, B., Latek, D., Yuan, S., Ghoshdastider, U., Debinski, A., & Filipek, S. (2012). Action of molecular switches in GPCRs - theoretical and experimental studies. Current Medicinal Chemistry, 19(8), 1090-1109. Retrieved from https://www.ncbi.nlm.nih.gov/pubmed/22300046

Vass, M., Podlewska, S., de Esch, I. J. P., Bojarski, A. J., Leurs, R., Kooistra, A. J., & de Graaf, C. (2019). Aminergic GPCR-Ligand Interactions: A Chemical and Structural Map of Receptor Mutation Data. Journal of Medicinal Chemistry, 62(8), 3784–3839. doi:10.1021/acs.jmedchem.8b00836

Vilardaga, J. P., Bunemann, M., Krasel, C., Castro, M., & Lohse, M. J. (2003). Measurement of the millisecond activation switch of G protein-coupled receptors in living cells. Nature Biotechnology, 21(7), 807–812. doi:10.1038/nbt838

Vogel, R., Mahalingam, M., Ludeke, S., Huber, T., Siebert, F., & Sakmar, T. P. (2008). Functional role of the “ionic lock” - an interhelical hydrogen-bond network in family A heptahelical receptors. Journal of Molecular Biology, 380(4), 648–655. doi:10.1016/j.jmb.2008.05.022

Wang, C., Jiang, Y., Ma, J. M., Wu, H. X., Wacker, D., Katritch, V., Han, G. W., Liu, W., Huang, X. P., Vardy, E., McCorvy, J. D., Gao, X., Zhou, X. E., Melcher, K., Zhang, C. H., Bai, F., Yang, H. Y., Yang, L. L., Jiang, H. L., Roth, B. L., Cherezov, V., Stevens, R. C., & Xu, H. E. (2013). Structural basis for molecular recognition at serotonin receptors. Science, 340(6132), 610–614. doi:10.1126/science.1232807

Wassenaar, T. A., Ingolfsson, H. I., Bockmann, R. A., Tieleman, D. P., & Marrink, S. J. (2015). Computational lipidomics with insane: a versatile tool for generating custom membranes for molecular simulations. Journal of Chemical Theory and Computation, 11(5), 2144–2155. doi:10.1021/acs.jctc.5b00209

Wassenaar, T. A., Pluhackova, K., Bockmann, R. A., Marrink, S. J., & Tieleman, D. P. (2014). Going backward: a flexible geometric approach to reverse transformation from coarse grained to atomistic models. Journal of Chemical Theory and Computation, 10(2), 676–690. doi:10.1021/ct400617g

Yin, W., Zhou, X. E., Yang, D., de Waal, P. W., Wang, M., Dai, A., Cai, X., Huang, C. Y., Liu, P., Wang, X., Yin, Y., Liu, B., Zhou, Y., Wang, J., Liu, H., Caffrey, M., Melcher, K., Xu, Y., Wang, M. W., Xu, H. E., & Jiang, Y. (2018). Crystal structure of the human 5-HT1B serotonin receptor bound to an inverse agonist. Cell Discovery, 4, 12. doi:10.1038/s41421-018-0009-2

Yuan, D., Liu, Z., Kaindl, J., Maeda, S., Zhao, J., Sun, X., Xu, J., Gmeiner, P., Wang, H. W., & Kobilka, B. K. (2020). Activation of the alpha2B adrenoceptor by the sedative sympatholytic dexmedetomidine. Nature Chemical Biology, 16(5), 507–512. doi:10.1038/s41589-020-0492-2

Zhu, K., Pincus, D. L., Zhao, S. W., & Friesner, R. A. (2006). Long loop prediction using the protein local optimization program. Proteins-Structure Function and Bioinformatics, 65(2), 438–452. doi:10.1002/prot.21040

