## Supplementary files for "Binding and activation of serotonergic G-protein coupled receptors by the multimodal antidepressant vortioxetine"

Supporting Figure S1: Comparison of the three GPCR models

Supporting Figure S2: Vortioxetine binding modes in 5-$HT_{1A}$ from IFD and post MD

Supporting Figure S3: Vortioxetine binding modes in 5-$HT_{1B}$ from IFD and post MD

Supporting Figure S4: Vortioxetine binding modes in 5-$HT_7$ from IFD and post MD

Supporting Figure S5: $H6_{IC}$-$H3_{IC}$ distance in short MD simulations

Supporting Figure S6: Intermolecular 3.32-VXT-7.39 coordination

Supporting Figure S7: Receptor activation in apo 5-$HT_{1A}$

Supporting Figure S8: Sequence alignment used in homology modelling.

Supporting Table S1: Percent identity matrix of sequence identities amongst serotonergic GPCRS

Supporting Table S2: Data from the docking calculations

Supporting Table S3: Relative free energy of binding of vortioxetine to each protein

Supporting Table S4: Non-conserved residues in the LBS in the serotonergic GPCRs

SUPPORTING FIGURES

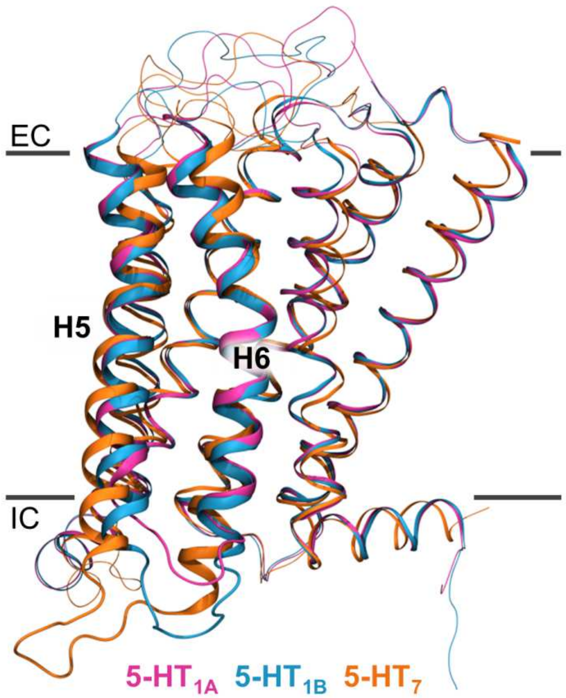


**Supporting Figure S1.** Comparison of the three GPCR models; 5-$HT_{1A}$ (pink), 5-$HT_{1B}$ (blue), and 5-$HT_7$ (orange). The 5-$HT_{1A}$ and 5-$HT_{1B}$ receptors are in active-like conformational states and the 5-$HT_7$ receptor is in an inactive conformational state as indicated by the differences in conformation of H5 and H6 shown in thicker ribbons. The conformation of $H6_{IC}$ is similar in all three models due to the lack of bound G-protein in the active-like models. The approximate location of the membrane is indicated and the extra- (EC) and intracellular (IC) side of the membrane is noted. The receptors are aligned on Cα atoms of H1-4.

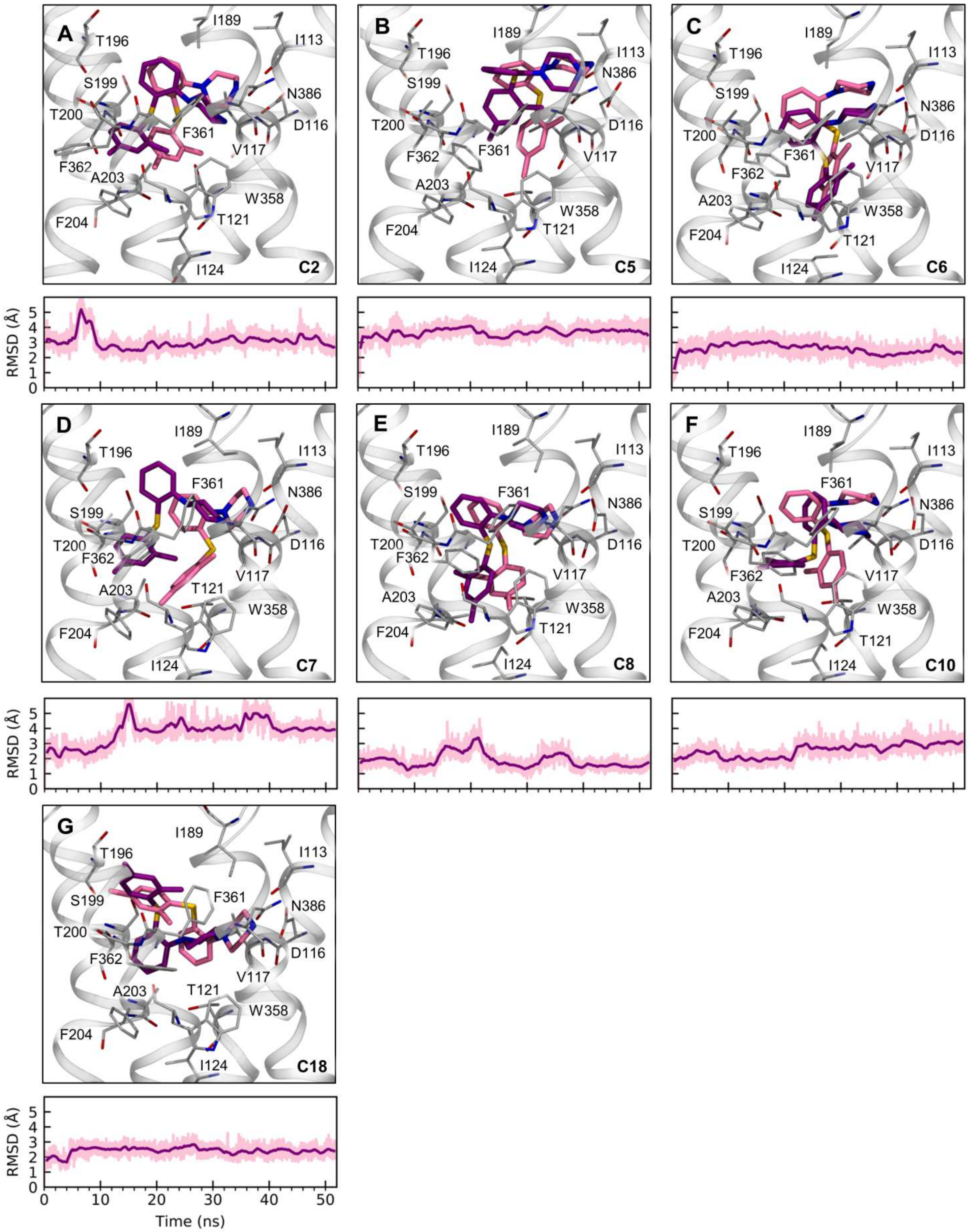


**Supporting figure S2.** Overlays of the first and last conformation of vortioxetine during MD simulated relaxation for a representative pose from binding cluster **A**) C2, **B**) C5, **C**) C6, **D**) C7, **E**)

C8, **F**) C10, and **G**) C18 found for vortioxetine binding to the 5-$HT_{1A}$ receptor. The lighter pink represents the conformation of vortioxetine as found in the IFD calculations, while the dark pink represents the same conformation of vortioxetine following 52 ns of MD simulation. Beneath each panel the time-progressed RMSD of vortioxetine heavy atoms is reported. The raw data is shown in light pink while the running-average is shown in dark pink.

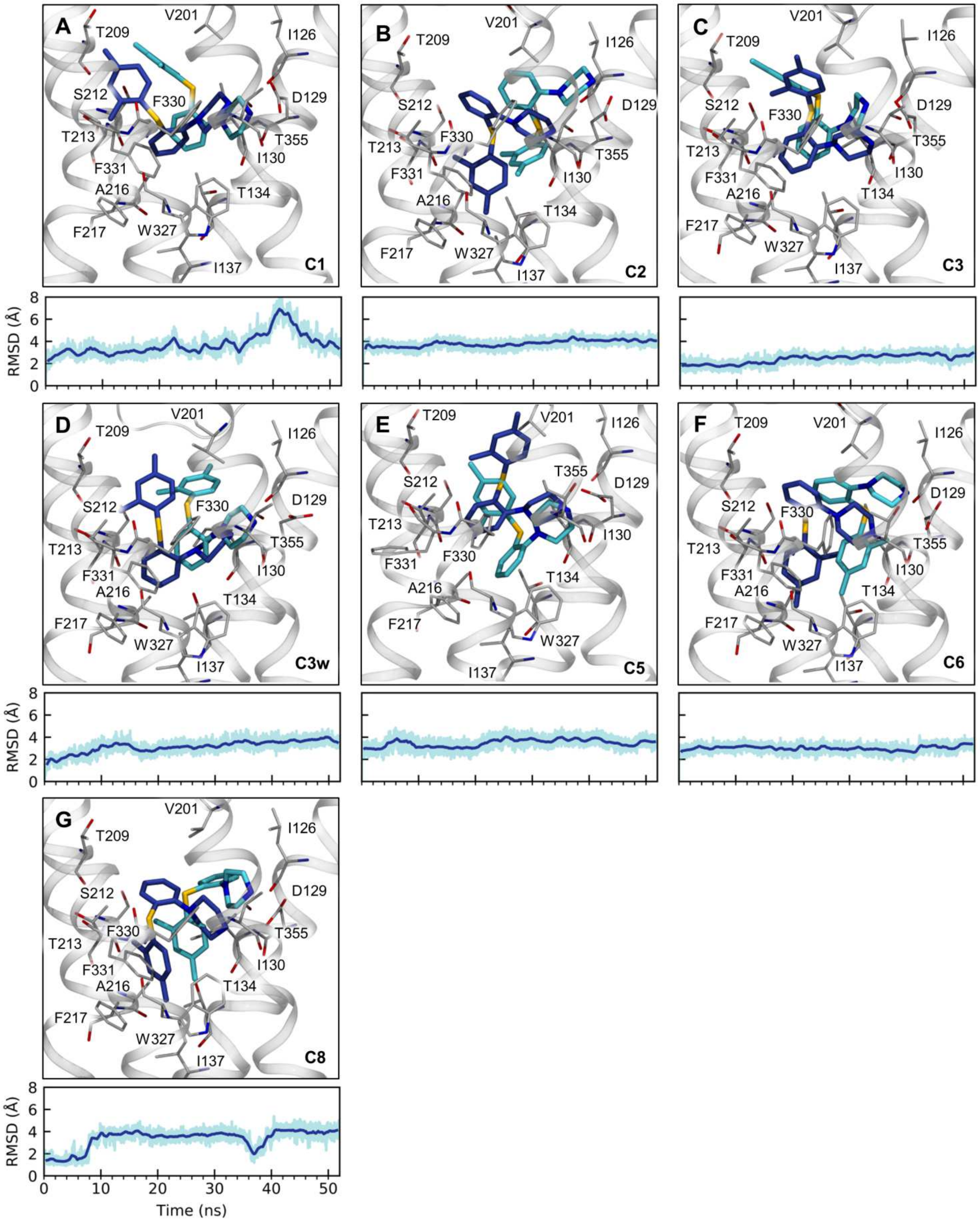


**Supporting figure S3**. Overlays of the first and last conformation of vortioxetine during MD simulated relaxation for a representative pose from binding cluster **A**) C1, **B**), C2 **C**), C3 **D**), C3w **E**)

C5, **F**) C6, and **G**) C8 found for vortioxetine binding to the 5-$HT_{1B}$ receptor. The lighter blue represents the conformation of vortioxetine as found in the IFD calculations, while the dark blue represents the same conformation of vortioxetine following 52 ns of MD simulation. Beneath each panel the time-progressed RMSD of vortioxetine heavy atoms is reported. The raw data is shown in light blue while the running-average is shown in dark blue.

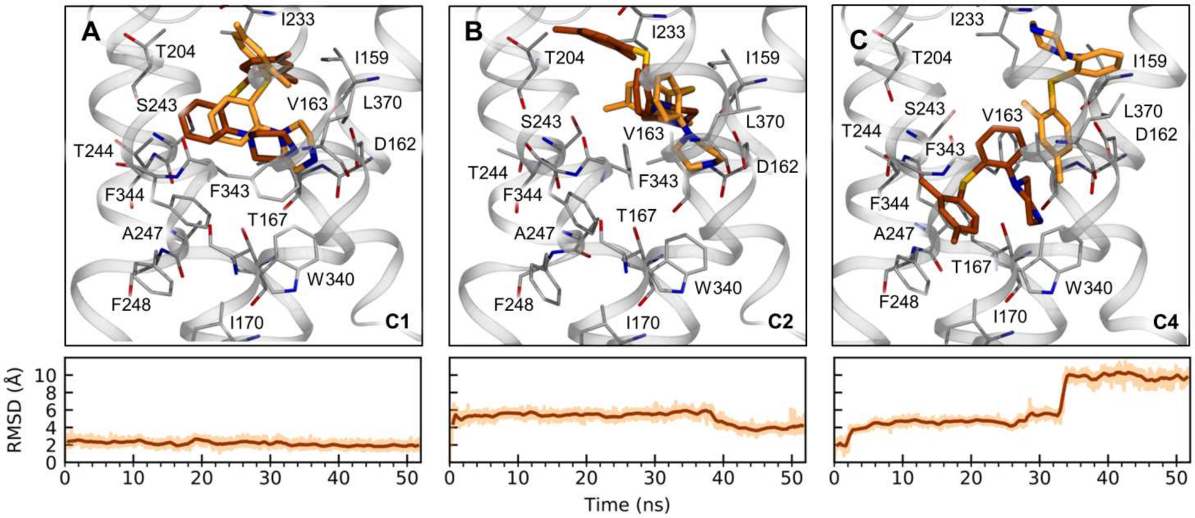


**Supporting figure S4**. Overlays of the first and last conformation of vortioxetine during MD simulated relaxation for a representative pose from binding cluster **A**) C1, **B**) C2, and **C**) C4 found for vortioxetine binding to the 5$HT_7$ receptor. The lighter orange represents the conformation of vortioxetine as found in the IFD calculations, while the dark orange represents the same conformation of vortioxetine following 52 ns of MD simulation. Beneath each panel the time-progressed RMSD of vortioxetine heavy atoms is reported. The raw data is shown in pale orange while the running-average is shown in dark orange.

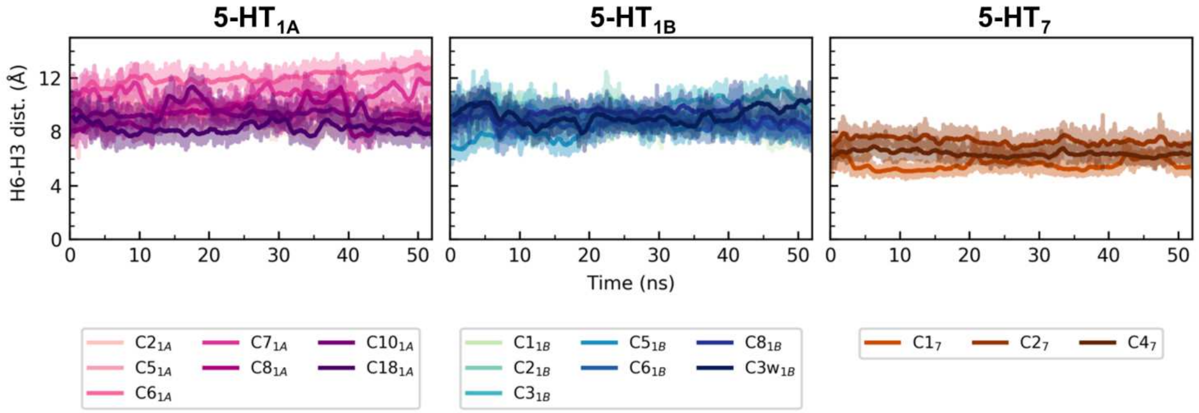


**Supporting Figure S5**. The distance between $H6_{IC}$-$H3_{IC}$ in the short MD simulations of receptor/vortioxetine complexes. The raw data is shown in transparent hues, while the running average is shown in opaque hues.

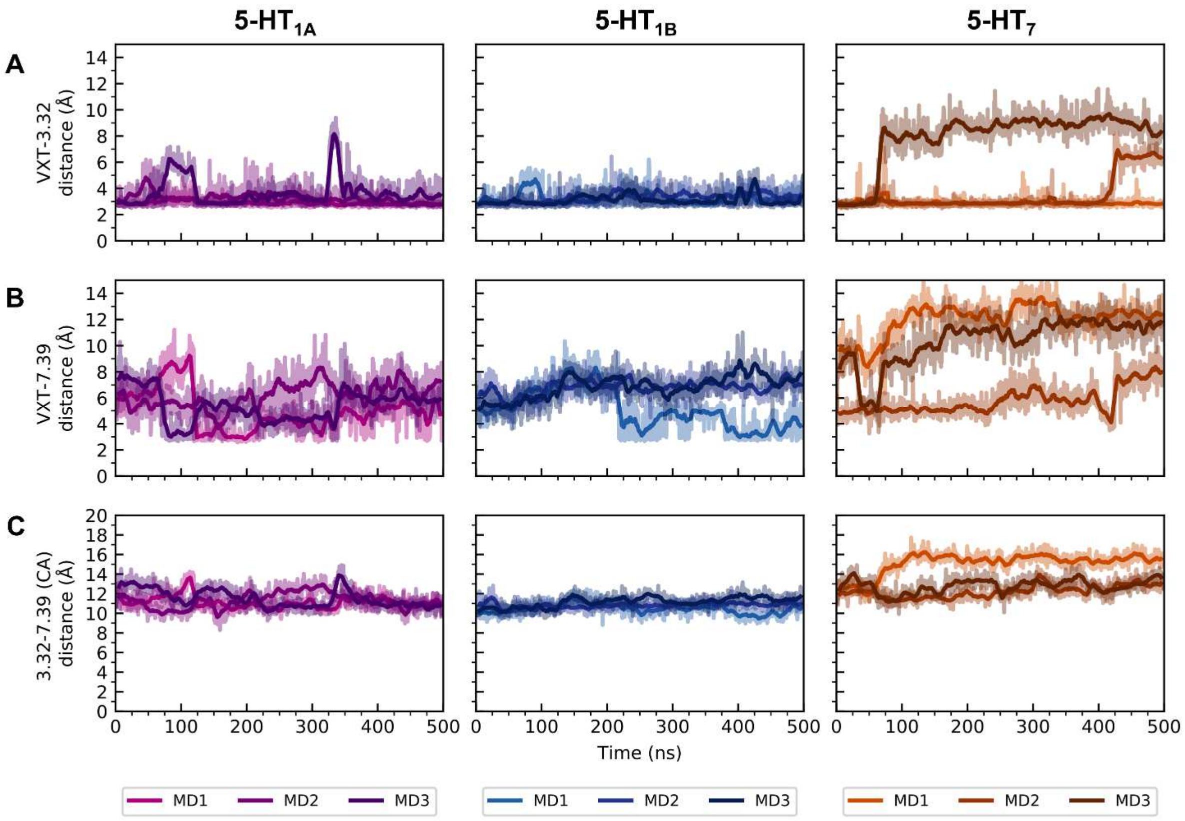


**Supporting Figure S6**. Intermolecular 3.32-VXT-7.39 coordination. **A**) The distance between the charged amine in vortioxetine and the charged Asp3.32. **B**) The distance between the charged amine in vortioxetine and the sidechain of Asn7.39 (5-$HT_{1A}$), Thr7.39 (5-$HT_{1B}$) or Leu7.39 (5-$HT_7$). The distances were calculated as the minimal distance between amine and either extreme sidechain atom. **C**) The Cα-Cα distance between Asp3.32 and Asn7.39 (5-$HT_{1A}$), Thr7.39 (5-$HT_{1B}$) or Leu7.39 (5-$HT_7$). The raw data is shown in transparent hues, while the running average is shown in opaque hues.

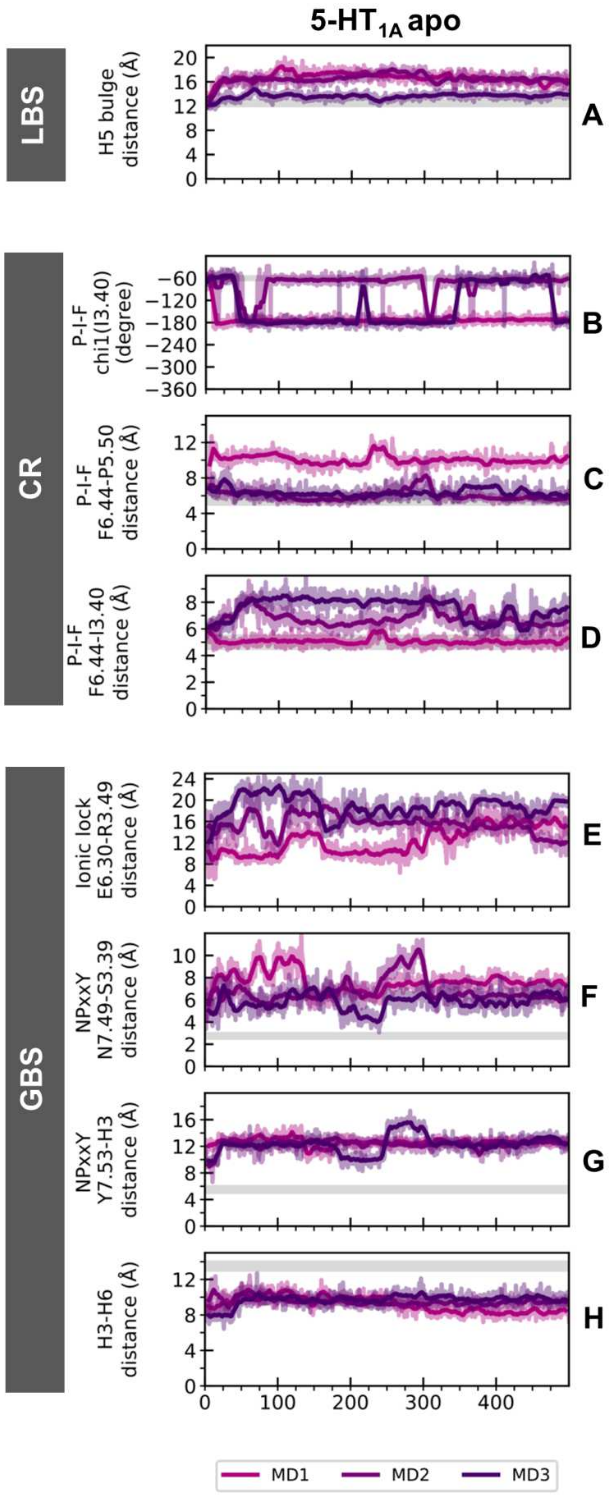


**Supporting Figure S7**. Measures of receptor activation for 5-$HT_{1A}$ in the apo state. The time-progressed degree or distance changes are shown for **A**) the LBS, **B-D**) the connector region (CR), and **E-H**) the GBS. In each panel, the active-state value of each measure is indicated by a horizontal, gray line, except for the ionic lock panel, which could take on any value above 4 Å when in an active state. For each panel, the raw data is shown in transparent and the smoothed data is shown in opaque hues. The details on how each parameter was calculated can be found in the Methods section.

**A**

```
h5-HT1B/1-275     1 YIYQDSISLPWKVLLVMLLALITLATTLSNAFVIATVYRTRKLHTPANYLIASLAVTDLLVSILVMPIST  70
h5-HT1A/1-275     1 TTGISDVTVSYQVITSLLLGTLIFCAVLGNACVVAAIALERSLQNVANYLIGSLAVTDLMVSVLVLPMAA  70

h5-HT1B/1-275    71 MYTVTGRWTLGQVVCDFWLSSDITCCTASIWHLCVIALDRYWAITDAVEYSAKRTPKRAAVMIALVWVFS 140
h5-HT1A/1-275    71 LYQVLNKWTLGQVTCDLFIALDVLCCTSSILHLCAIALDRYWAITDPIDYVNKRTPRRAAALISLTWLIG 140

h5-HT1B/1-275   141 ISISLPPFFWRQAKAEEEVSECVVNTDHILYTVYSTVGAFYFPTLLLIALYGRIYRERKATKTLGIILGA 210
h5-HT1A/1-275   141 FLISIPPMLGWRTPEDRSDPDACTISKDHGYTIYSTFGAFYIPLLLMLVLYGRIFRERKTVKTLGIIMGT 210

h5-HT1B/1-275   211 FIVCWLPFFIISLVMPICKDACWFHLAIFDFFTWLGYLNSLINPIIYTMSNEDFKQAFHKLIRFK      275
h5-HT1A/1-275   211 FILCWLPFFIVALVLPFCESSCHMPTLLGAIINWLGYSNSLLNPVIYAYFNKDFQNAFKKIIKCK      275
```

**B**

```
hD3/32-400       32 --------YALSYCALILAIVFGNGLVCMAVLKERALQTTTNYLVVSLAVADLLVATLVMPWVVYLEVTG  93
h5-HT7/1-285      1 YGRVEKVVIGSILTLITLLTIAGNCLVVISVCFVKKLRQPSNYLIVSLALADLSVAVAVMPFVSVTDLIG  70

hD3/32-400       94 GVWNFSRICCDVFVTLDVMMCTASIWNLCAISIDRYTAVVMPVHYQHGTGQSSCRRVALMITAVWVLAFA 163
h5-HT7/1-285     71 GKWIFGHFFCNVFIAMDVMCCTASIMTLCVISIDRYLGITRPLTYP---VRQNGKCMAKMILSVWLLSAS 137

hD3/32-400      164 VSCPLLFGF-NTTGDPTVCSIS-NPDFVIYSSVVSFYLPFGVTVLVYARIYVVLKQRRRK---GVPLREK 228
h5-HT7/1-285    138 ITLPPLFGWAQNVNDDKVCLISQDFGYTIYSTAVAFYIPMSVMLFMYYQIYKAARKSAAKKNISIFKREQ 207

hD3/32-400      229 KATQMVAIVLGAFIVCWLPFFLTHVLNTHC---QTCHVSPELYSATTWLGYVNSALNPVIYTTFNIEFRK 295
h5-HT7/1-285    208 KAATTLGIIVGAFTVCWLPFFLLSTARPFICGTSCSCIPLWVERTFLWLGYANSLINPFIYAFFNRDLRT 277

hD3/32-400      296 AFLKILSC                                                               303
h5-HT7/1-285    278 TYRSLLQC                                                               285
```

**Supporting Figure S8**. Sequence alignments of the **A**) 5-$HT_{1A}$ and 5-$HT_{1B}$ and **B**) 5-$HT_7$ and D3 receptor sequences used in the homology modeling. Only the modeled parts of the sequences are included in the alignment, and the residue numbering is therefore not comparable to the canonical sequences. The underlying uniprot/PDB IDs are: P08908, 4IAR, P34969, and 3PBL, respectively.

SUPPORTING TABLES

**Supporting Table S1**. Percent identity matrix of all serotonergic GPCRs except for 5-$HT_{5B}$. The subtypes of the 5-$HT_1$ and 5-$HT_2$ receptors are boxed in. Based on alignment by Clustal Ω 2.1.

| | **1A** | **1B** | **1D** | **1E** | **1F** | **2A** | **2B** | **2C** | **4** | **5A** | **6** | **7** |
|---|---|---|---|---|---|---|---|---|---|---|---|---|
| **1A** | 100 | 44 | 43 | 40 | 41 | 29 | 30 | 31 | 40 | 34 | 32 | 38 |
| **1B** | 44 | 100 | 62 | 47 | 49 | 28 | 25 | 28 | 35 | 35 | 30 | 38 |
| **1D** | 43 | 62 | 100 | 48 | 48 | 30 | 27 | 30 | 34 | 34 | 31 | 36 |
| **1E** | 40 | 47 | 48 | 100 | 56 | 32 | 28 | 31 | 35 | 33 | 30 | 37 |
| **1F** | 41 | 49 | 48 | 56 | 100 | 30 | 26 | 30 | 37 | 35 | 29 | 37 |
| **2A** | 29 | 28 | 30 | 32 | 30 | 100 | 46 | 52 | 30 | 27 | 28 | 28 |
| **2B** | 30 | 25 | 27 | 28 | 26 | 46 | 100 | 45 | 28 | 26 | 27 | 25 |
| **2C** | 31 | 28 | 30 | 31 | 30 | 52 | 45 | 100 | 32 | 28 | 28 | 25 |
| **4** | 40 | 35 | 34 | 35 | 37 | 30 | 28 | 32 | 100 | 30 | 30 | 36 |
| **5A** | 34 | 35 | 34 | 33 | 35 | 27 | 26 | 28 | 30 | 100 | 31 | 35 |
| **6** | 32 | 30 | 31 | 30 | 29 | 28 | 27 | 28 | 30 | 31 | 100 | 32 |
| **7** | 38 | 38 | 36 | 37 | 37 | 28 | 25 | 25 | 36 | 35 | 32 | 100 |

**Supporting Table S2.** Results from the IFD calculations of vortioxetine into the 5-$HT_{1A}$, 5-$HT_{1B}$, and 5-$HT_7$ receptor. Average Emodel, Gscore, and IFD scores (kcal/mol) with accompanying standard deviations and number of poses within the cluster are reported for each binding cluster. The number of outliers in each calculation is also reported.

**5-$HT_{1A}$**

| Cluster | Poses | Av. Emodel | Av. Gscore | Av. IFDscore |
|---|---|---|---|---|
| $C2_{1A}$ | 2 | -43.6 ±3.9 | -8.2 ±0.2 | -511.1 ±0.4 |
| $C5_{1A}$ | 7 | -53.8 ±4.8 | -7.4 ±1.1 | -510.6 ±1.3 |
| $C6_{1A}$ | 17 | -49.1 ±4.0 | -6.9 ±0.8 | -510.0 ±0.9 |
| $C7_{1A}$ | 2 | -59.3 ±3.2 | -10.0 ±0.2 | -512.7 ±0.1 |
| $C8_{1A}$ | 28 | -53.7 ±7.3 | -8.5 ±0-9 | -511.5 ±1.0 |
| $C10_{1A}$ | 15 | -51.0 ±6.7 | 7.3 ±0.8 | -510.5 ±1.0 |
| $C18_{1A}$ | 5 | -48.8 ±13 | -7.2 ±1.0 | -510.6 ±1.3 |
| Outliers | 22 | - | - | - |

**5-$HT_{1B}$**

| Cluster | Poses | Av. Emodel | Av. Gscore | Av. IFDscore |
|---|---|---|---|---|
| $C1_{1B}$ | 26 | -53.1 ±8.4 | -7.8 ±0.8 | -569.1 ±0.9 |
| $C2_{1B}$ | 10 | -53.2 ±8.5 | -7.6 ±1.0 | -568.9 ±1.0 |
| $C3_{1B}$ | 9 | -54.4 ±9.0 | -7.9 ±1.1 | -569.1 ±1.3 |
| $C5_{1B}$ | 6 | -54.7 ±7.8 | -8.4 ±0.6 | -569.5 ±0.8 |
| $C6_{1B}$ | 4 | -53.4 ±3.9 | -8.4 ±0.3 | -569.5 ±0.5 |
| $C8_{1B}$ | 2 | -53.5 ±4.0 | -8.5 ±1.5 | -569.4 ±1.6 |
| $C3w_{1B}$ | 8 | -59.1 ±8.4 | -8.9 ±0.9 | -571.4 ±1.0 |
| Outliers | 40 | - | - | - |

**5-$HT_7$**

| Cluster | Poses | Av. Emodel | Av. Gscore | Av. IFDscore |
|---|---|---|---|---|
| $C1_7$ | 12 | -51.8 ±9.5 | -8.5 ±1.0 | -548.7 ±1.1 |
| $C2_7$ | 11 | -54.7 ±4.9 | -8.6 ±0.9 | -548.5 ±1.1 |
| $C4_7$ | 9 | -55.1 ±8.1 | -7.3 ±1.0 | -547.1 ±1.0 |

| Outliers | 9 | - | - | - |
|---|---|---|---|---|

**Supporting Table S3.** Average estimates of the relative free energy of binding of vortioxetine to each protein based on the MM-PBSA approach. Binding free energies were calculated based on 100 snapshots evenly extracted from the first 2 ns of each simulation as described in Methods. As the energies are relative it is not meaningful to compare their values across receptors. All energies are reported in kcal/mol relative to the least favorable free energy along with standard deviations for each binding cluster.

| 5-$HT_{1A}$ | | 5-$HT_{1B}$ | | 5-$HT_7$ | |
|---|---|---|---|---|---|
| Cluster | $\Delta\Delta G_{bind}$ | Cluster | $\Delta\Delta G_{bind}$ | Cluster | $\Delta\Delta G_{bind}$ |
| $C2_{1A}$ | 0 | $C1_{1B}$ | -8.6 ±1.0 | $C1_7$ | -29.5 ±1.3 |
| $C5_{1A}$ | -6.1 ±1.9 | $C2_{1B}$ | -20.2 ±0.9 | $C2_7$ | -27.5 ±1.5 |
| $C6_{1A}$ | -9.8 ±1.7 | $C3_{1B}$ | -9.9 ±0.9 | $C4_7$ | 0 |
| $C7_{1A}$ | -11.0 ±1.7 | $C5_{1B}$ | 0 | | |
| $C8_{1A}$ | -8.6 ±1.7 | $C6_{1B}$ | -17.1 ±0.9 | | |
| $C10_{1A}$ | -6.7 ±1.7 | $C8_{1B}$ | -16.5 ±0.8 | | |
| $C18_{1A}$ | -1.3 ±1.7 | $C3w_{1B}$ | -3.1 ±1.0 | | |

**Supporting Table S4**: Non-conserved residues in the LBS of the serotonergic GPCRs. The underlying multiple sequence alignment includes all serotonergic receptors (Clustal Ω 2.1).

| | 1A | 1B | 1D | 1E | 1F | 2A | 2B | 2C | 4 | 5A | 6 | 7 |
|---|---|---|---|---|---|---|---|---|---|---|---|---|
| **3.28** | F | W | W | W | W | W | W | W | R | W | W | F |
| **3.29** | I | L | L | L | L | I | L | I | T | I | T | I |
| **3.33** | V | I | I | M | I | V | V | V | V | V | V | V |
| **5.39** | T | T | T | T | T | V | M | V | A | A | V | T |
| **5.42** | S | S | S | S | S | G | G | G | C | S | A | S |
| **5.43** | T | T | T | T | T | S | S | S | S | T | S | T |
| **5.46** | A | A | A | A | A | S | A | A | A | A | T | A |
| **6.55** | A | S | S | E | E | N | N | N | N | E | N | S |
| **7.39** | N | T | T | T | A | V | V | V | L | L | T | L |
